# Inorganic pyrophosphate disrupts amorphous hydrated bone mineral interfaces in hypophosphatasia

**DOI:** 10.1101/2025.11.24.689921

**Authors:** Scott Dillon, Amelia Armiger, Adrian Murgoci, Linda Skingle, Fabiana G.A. Tabegna, Sarah McDonald, Steven Mumm, Michael P. Whyte, Mark Garton, Kenneth E.S. Poole, Melinda J. Duer

## Abstract

Bone mineral molecular architecture is tightly regulated by the kinetics of calcium phosphate phase transformations. In the rare skeletal disease hypophosphatasia (HPP), caused by inactivating mutations in the *ALPL* gene encoding tissue-nonspecific alkaline phosphatase (TNSALP), accumulation of inorganic pyrophosphate (PPi) alters these phase dynamics. Using solid-state nuclear magnetic resonance spectroscopy and high-resolution electron microscopy, in patient samples we show that bone mineral from compound heterozygous HPP patients exhibits a loss of hydrated amorphous interfacial phases and instead contains highly crystalline hydroxyapatite (HAP), correlating with abnormally high bone mineral density and brittle atypical femoral fractures. Synthetic and cellular models demonstrate that elevated PPi impedes normal phase transitions from amorphous calcium phosphate precursors, bypassing intermediate states and driving ordered HAP nucleation. These findings link disrupted mineral phase kinetics to pathological bone mineral molecular structure, emphasising the critical importance of the hydrated amorphous shell around bone mineral for its material properties and redefining HPP as a molecular mineralization disorder. Our findings also illustrate how biochemical cues regulate non-equilibrium crystallization pathways in biomineralization.

## Introduction

Bone tissue is a biological composite material of inorganic calcium phosphate mineral embedded within and around extracellular organic collagen I protein fibrils(*1*). Both its composite nature and the hierarchical structures which mineralised collagen fibrils are built into in biological tissues convey remarkable material properties, allowing both flexibility and strength under mechanical stress at length scales spanning individual fibrils to a whole organ(*2*, *3*). The molecular structure of the mineral phase has been the focus of decades of interdisciplinary research, revealing a thin poorly crystalline carbonated and substituted hydroxyapatite (HAP) platelet core, surrounded by amorphous, highly hydrated non-apatitic material (*4–7*). High resolution electron microscopy studies show that these platelets are arranged in stacks within and around collagen fibrils, making the amorphous non-apatitic material the interface between HAP platelets(*1*, *8–11*). These individual platelet units are bridged by large metabolite anions including citrate embedded in the hydrated envelope, which separate them and prevent adjacent units collapsing and transforming to the thermodynamically favourable crystalline mineral(*12*, *13*).

However, the mechanisms employed by mineralising cells to manipulate the physical chemistry governing phase dynamics required to generate this intricate mineral molecular architecture remain unclear, and the importance of the interfacial amorphous material remains largely unexplored. Calcium phosphate phase dynamics are complex and can result in the formation of a wide range of materials in solution, often through transient metastable states, depending on pH, temperature, pressure, ionic strength and other factors such as confinement(*14–17*). These physiochemical dynamics are likely complicated further in biological systems by cellular regulation, interaction with biomolecules, and spatial and temporal confinement imposed by the organic matrix(*11*, *18*, *19*). Pathological conditions in which biomineralisation is impaired, such as those arising from mutations in proteins essential for bone calcification, therefore provide a valuable framework for probing these mechanisms.

The rare bone disease hypophosphatasia (HPP) is caused by inactivating mutation(s) in the gene *ALPL* which encodes the tissue-nonspecific isoenzyme of alkaline phosphatase (TNSALP)(*20*). TNSALP is a homodimeric ectoenzyme present on the external surface of mineralizing cells which cleaves several phosphomonoester substrates including inorganic pyrophosphate (PPi) to release inorganic phosphate(*21*). PPi is thought to act as a potent mineralization inhibitor by binding to nascent mineral surfaces and preventing crystal growth(*22*, *23*). The balance between inhibition of crystal growth and the controlled provision of crystal building blocks regulated by TNSALP is therefore thought to provide some control over physiochemical phase transitions during bone mineral nucleation.

In HPP, accumulation of PPi results in a wide range of clinical symptoms, which can include bone pain, weakness and expansion of unmineralised material (osteoid) associated with pseudofractures at the metatarsals, known as osteomalacia (OM) in adults(*24*, *25*). A severe subcohort of HPP patients carry compound heterozygous *ALPL* mutations with very low residual enzymatic activity(*26*). These patients often present in later adulthood with bilateral subtrochanteric brittle stress fractures, similar to atypical femoral fractures (AFFs) observed as a complication of long-term bisphosphonate treatment for low bone mass conditions(*26–28*). These fractures are distinct from the osteomalacic pseudofractures more commonly associated with HPP. The occurrence of spontaneous complete fractures at bilateral sites of maximal predicted femoral tensile force in such patients remains an enigma.

The elevated PPi in HPP also explains the disorder’s chondrocalcinosis from calcium pyrophosphate dihydrate deposition(*29*), pseudogout as a PPi arthropathy, and enigmatic calcific periarthritis featuring inflammation from soft tissue accumulation of HAP wherein elevated PPi can promote HAP deposition(*30*, *31*). Excess PPi might explain the HPP complication of chronic recurrent multifocal osteomyelitis(*32*, *33*). Rarely in adults with HPP there can be enigmatic paradoxical progressive elevation of BMD(*34*) and in affected children patchy areas of osteosclerosis(*35*).

Here we use solid state nuclear magnetic resonance (SSNMR) spectroscopy in combination with high resolution electron microscopy to show that bone mineral molecular architecture is radically altered in compound heterozygous HPP patients and results in highly crystalline HAP in which the hydrated non-apatitic interfaces between HAP platelets are disrupted, causing high bone mineral density (BMD) and brittle atypical fractures. Our data explicitly link the clinical phenotype in patients with their bone mineral chemistry for the first time and expose the physical chemistry underlying the pathophysiology of atypical fractures in HPP. We further demonstrate that PPi regulates the kinetics of calcium phosphate phase transformations and that these disrupted physiochemical kinetics are responsible for aberrant mineral molecular architecture in HPP.

### Compound heterozygous HPP patients present with bilateral atypical femoral fractures, high vertebral bone mineral density and hyperosteoidosis

Sanger sequencing (*36*) identified three patients from the Eastern Region Rare Bone clinic, (ErBON), part of the NHS Rare Diseases Collaborative Network (RDCN) at Addenbrooke’s hospital, Cambridge, UK, carrying compound heterozygous mutations in *ALPL*, two of whom are reported here. Patient 0124 (49 year-old Caucasian female) exhibited a novel missense variant (c.536C>G, p.Ala179Gly), and a rare nonsense defect (c.891C>A, p.Tyr297*), retaining a predicted 2.8% enzyme function(*37*). The missense variant was predicted to be probably damaging with a score of 0.973 by Polymorphism Phenotyping v2 (PolyPhen-2)(*38*), and was predicted to affect protein function with a score of 0.02 by Sorting Intolerant from Tolerant (SIFT)(*39*). *ALPL* Ala179Gly is extremely rare (0.000186%) in a healthy population (gnomAD: https://gnomad.broadinstitute.org). Another *ALPL* variant (Ala179Thr) at the same amino acid position is pathogenic. These factors, combined with presence of a pathogenic variant (Tyr297*) on the patient’s other allele make Ala179Gly “Likely pathogenic” according to the American College of Medical Genetics (ACMG) guidelines(*40*). Patient 1716 (54 year-old Caucasian female) exhibited compound heterozygous missense mutations, namely c.526G>A (p.Ala176Thr; 45.4% predicted enzyme activity)(*41*) and c.1171C>T (p.Arg391Cys; 2.2% predicted enzyme activity)(*37*). The three previously identified mutations in both patients are categorised as pathogenic in the *ALPL* Gene Variant Database(*42*). Both patients demonstrated consistently low serum alkaline phosphate of <5 U/L.

Both patients presented with atraumatic complete subtrochanteric AFFs of the right femur and contralateral subtrochanteric pseudofractures with no history of bisphosphonate treatment. Other radiological findings in patient 1716 have been previously reported elsewhere(*43*). Areal bone mineral density (aBMD) measured using dual X-ray absorptiometry (DXA) in the lumbar spine revealed extremely high BMD (Z(T) patient 0124: +6.6(+5.9); patient 1716: +2.5(+3.6)). This was confirmed using quantitative computed tomography (qCT) to measure trabecular volumetric BMD (vBMD) at L3, revealing densities of 374 mg/cm^3^ (Z-score +9.04; T-score +7.77) in patient 0124 and 272 mg/cm^3^ (Z-score +5.77) in patient 1716 (Supp. Fig. S1).

Undecalcified transiliac crest bone biopsy histopathology was carried out to investigate abnormalities in bone mineralization and microarchitecture characteristic of HPP and were compared to those from three patients with other forms of osteomalacia (OM) (one with tumor-induced osteomalacia and two with X-linked hypophosphatemia; patients 0120, 0718 and 1816 respectively), as well as to three reference patients who were found to have normal mineralization (patients 0517, 0817 and 1517). Von Kossa staining revealed the expected marked increase in osteoid surface normalised to bone surface (OS/BS) in both HPP and OM patients compared to controls, however trabecular number (TbN) was also increased and separation (TbSp) reduced in HPP patients alone, indicating trabecular sclerosis (Supp. Fig. S1; Supp. Table 1).

### HPP bone mineral nanostructure is disrupted and is distinct from other forms of osteomalacia

We hypothesized that the increased fracture propensity and the atypical fracture patterns observed in compound heterozygous HPP patients arise from alterations in one or more of: the spatial distribution of bone mineral, the mineral composition, or the morphology of the constituent mineral nanoparticles that collectively make up bone mineral (Fig. 1, A). To investigate this hypothesis, we first compared the spatial distribution of bone mineral in trans-iliac crest biopsies obtained from patients with HPP, OM, and unaffected controls. Backscattered electron scanning electron microscopy (BSE-SEM) was employed to visualize the spatial location of mineral within bone trabeculae on the micron lengthscale (Fig. 1, B-D).

**Fig. 1.**
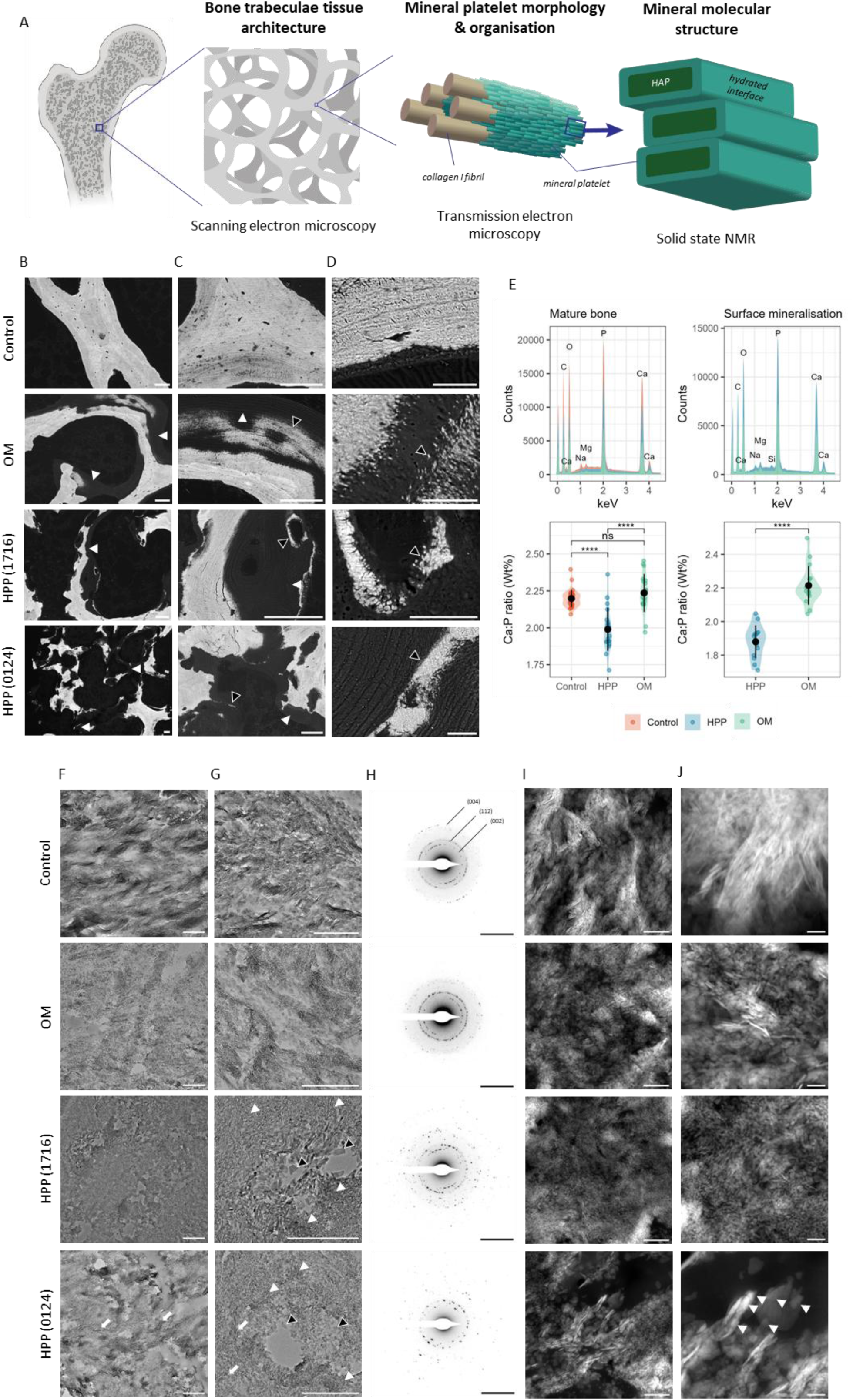
Nanostructural analysis of hypophosphatasia patient bone biopsies. **(A)** Schematic illustration of bone tissue structural hierarchy and techniques used to study different length scales. **(B-D)** Basckscattered electron scanning electron microscopy of bone biopsies. White arrowheads indicate regions of osteoid accumulation. Black arrowheads indicate surface mineralization. Scale bars = 100µm (B,C); 20µm (D). **(E)** Energy dispersive X-ray spectroscopy spectra and quantification of Ca:P wt % ratio. ****: p < 0.001. **(F,G)** Brightfield transmission electron microscopy in ultrathin biopsy sections. White arrowheads indicate regions containing small compacted crystals. Black arrowheads indicate large individual particles. White arrows indicate areas maintaining limited platelet stacking in hypophosphatasia patient 0124. Scale bars = 500nm. **(H)** Selected area electron diffraction. Scale bars = 5nm^-1^. **(I,J)** Scanning-transmission electron microscopy in ultrathin sections. White arrowheads indicate large crystals in hypophosphatasia comprised of smaller rounded crystallites. Scale bars = 200nm (I); 50nm (J).

Trabecular bone surfaces in control patients appeared smooth and quiescent as opposed to OM and HPP bone surfaces which appeared rough and scalloped (Fig. 1, B-D). Both OM and HPP patient samples exhibited accumulation of unmineralised osteoid on trabecular bone surfaces (Fig. 1, B,C; white arrowheads). Patchy mineralisation was observed within the osteoid seam in both OM and HPP samples, however in HPP samples this occurred only remote from the bone surface (Fig. 1, B,C; black arrowheads). In OM specimens this surface mineralisation occurred as nanoscopic mineralisation foci, as expected in physiological bone mineralisation, whereas in HPP samples it had a dense popcorn morphology (Fig. 1, D; black arrowheads). No differences were observed within the control patient group or between TIO and XLH patients. We concluded that HPP bone contains a distinctly different mineral spatial distribution to both typical OM and normal bone, and what appears to be forming mineral at bone surfaces has a significantly different microstructure to that in OM patients.

We next investigated the composition of the mineral phase using energy dispersive X-ray spectroscopy (EDS) to probe the Ca and P content in different spatial locations of the bone samples, quantifying the Ca:P ratio in point scan spectra for both bulk trabeculae and surface-associated mineral. A significant reduction in mean Ca:P ratio in HPP samples was observed compared to both control and OM specimens in trabecular bone (p < 0.001) and between HPP and OM samples at surface-associated mineralization (p < 0.001), indicating the presence of a distinct calcium phosphate phase to that in normal bone mineral (Fig. 1, E).

To probe the mineral nanoparticle morphology and how they are organized relative to each other in HPP compared to controls and OM bone, we then performed transmission electron microscopy (TEM) in ultrathin sections of patient bone biopsies. Brightfield TEM images showed stacks of thin nanoscopic mineral platelets in both control and OM tissue consistent with previous analyses of bone mineral nanostructure(*1*, *8*). HPP patient samples however exhibited regions of dense compacted crystallites without stacked platelet morphology (Fig. 1, F,G; white arrowheads) and contiguous regions of large (50 - 75 nm in width) individual crystals surrounding void spaces (Fig. 1 F,G; black arrowheads). HPP patient 0124 was observed to have a less severe nanostructural defect, with some areas exhibiting limited platelet stacking (Fig. 1, F,G; white arrows). Selected area electron diffraction (SAED) in large individual particles demonstrated reflections corresponding to HAP crystal planes with small lattice d-spacings, in addition to the strong reflections corresponding to the (002), (112) and (004) planes observed in control and OM platelet stacks (Fig. 1, H; Supp. Fig. S2; Supp. Table 1). This indicates an increase in crystallinity and long-range crystal order in HPP bone mineral compared to control and OM patients, but with a loss of normal bone nanostructural organisation.

The lack of stacked platelet mineral morphology in HPP bone suggests disruption to the hydrated mineral platelet envelope which separates adjacent platelets. We investigated this hypothesis using high-resolution scanning-transmission electron microscopy (STEM) with a high-angle annular darkfield (HAADF) detector to visualise crystal interfaces. In control and OM samples crystals appeared as thin single platelets 5 - 7 nm in width, separated by 2 - 4 nm electron lucent interfaces (Fig 1, I,J), from the expected interfaces between the hydrated envelopes on neighbouring mineral platelets (Fig. 1, A). In contrast, the dense compacted mineral regions in HPP samples appeared comprised of rounded particles which varied in size between 5 - 10 nm and which were separated by 1 – 2 nm interfaces. In regions with large individual particles, high resolution STEM images revealed these were also composed of smaller rounded crystallites (Fig. 1, J; white arrowheads).

### HPP bone mineral molecular structure exhibits loss of structural water and non-crystalline interfacial phases

We concluded from these EM studies that HPP bone mineral has a significantly different composition and nanostructure (mineral particle morphology and organization) to normal bone mineral. This suggests that HPP bone mineral contains different mineral phase/s to normal bone. To characterise the mineral phase in HPP bone and compare it to that in normal and OM bone, we used solid state NMR spectroscopy (ssNMR) in bone biopsy serial sections from HPP, OM and control patients to probe the detailed molecular structure of the mineral phase.

1D ^31^P{^1^H} cross polarisation (CP) spectra of bone typically exhibit overlapping, unresolved signals from the (many) different mineral phosphatic environments present in a sample, resulting in a single broad ^31^P NMR peak at δ(^31^P) ∼ 3 ppm, which we observe in all samples (Fig. 2, A). The linewidth of this ^31^P signal is a measure of the range of different phosphate environments in the mineral and is compared in Fig. 2, B for all samples. CP ssNMR spectra can be recorded as a function of CP contact time and spectra recorded with different contact times have a bias on which phosphate ^31^P signals have stronger intensity. ^31^P{^1^H} CP ssNMR spectra recorded with a long contact time (10 ms) exhibit signals from the orthophosphate ^31^P sites in

**Fig. 2.**
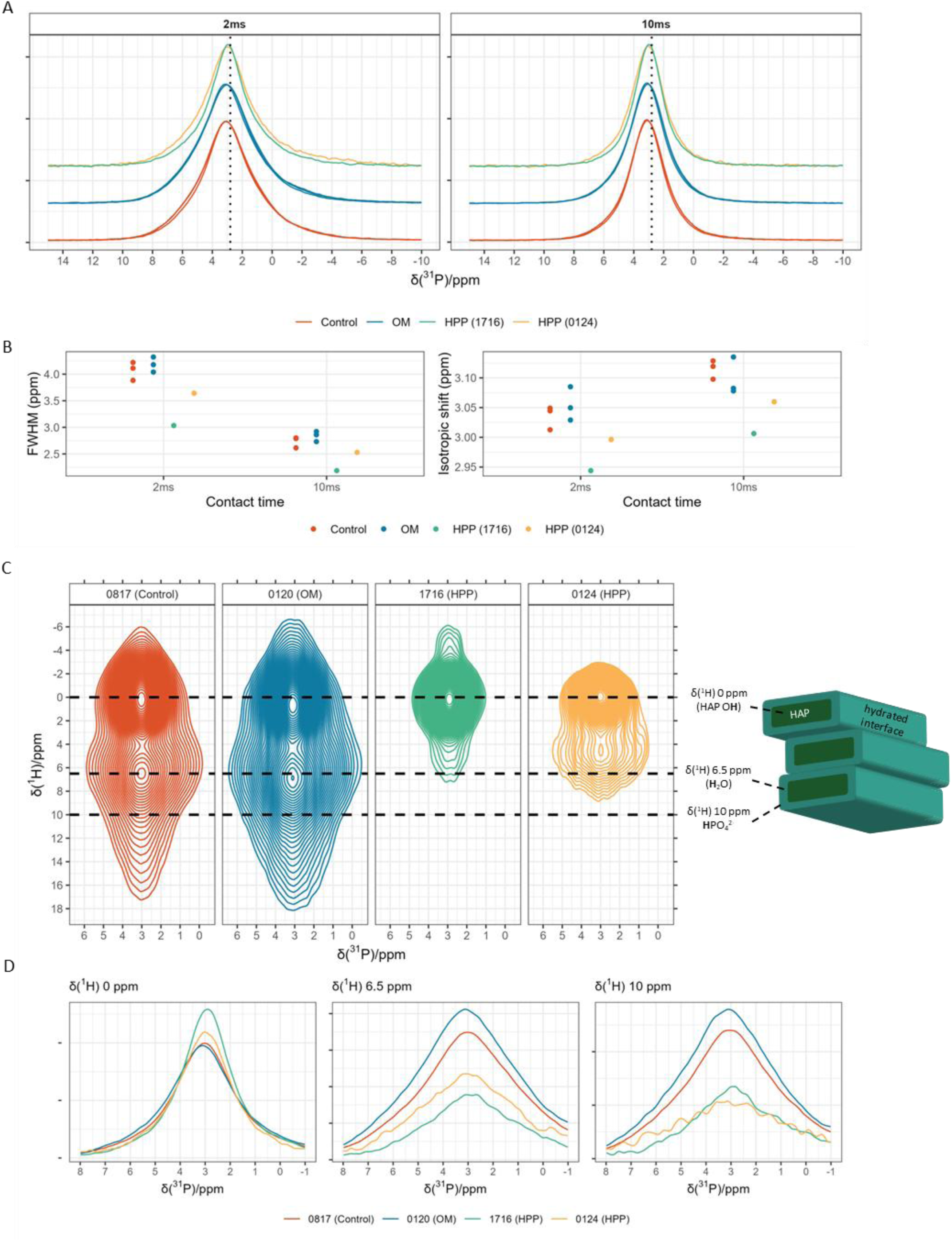
Molecular structure of hypophosphatasia bone by solid state nuclear magnetic resonance spectroscopy. **(A)** 1D ^31^P{^1^H} cross polarisation spectra at different contact times. **(B)** Full-width at half-maximum (FWHM) and isotropic peak position of spectra in (A), measured by fitting a Lorentzian function to each spectrum. **(C)** ^1^H-^31^P frequency-switched Lee-Goldburg heteronuclear correlation spectra representative of each patient group. ^1^H environment spectral assignments to the bone mineral molecular structure are indicated in the schematic on the right. Data are displayed with 40 contours between 20 – 100% normalised signal intensity. **(D)** 1D slices through spectra In (C) at dashed lines in the direct ^31^P dimension.

HAP almost exclusively, i.e. spectra at this long contact time are represent primarily the material at the core of mineral particles(*4*). The ^31^P signals in long contact time ^31^P{^1^H} CP NMR spectra of HPP bone have smaller linewidths compared to control and OM specimens, indicating higher mineral molecular order in the HAP mineral particle component over that for normal (and OM) bone mineral (Fig. 2, A,B). We also observe that the peak maxima in the HPP spectra are at lower ^31^P chemical shifts (2.9±0.03 ppm) compared to that for OM (3.1±0.03 ppm) and control (3.1±0.02 ppm) samples. The ^31^P peak maxima chemical shift depends on the size of the HAP domain and this reduction in chemical shift is consistent with the dominant phosphate environment in HPP bone mineral particles being more similar to that in large (microcrystalline or larger) HAP crystals, than the nanocrystalline HAP characteristic of bone mineral(*6*).

We then investigated the molecular structure of the interfacial hydrated envelopes of bone mineral particles using 2D ^1^H-^31^P heteronuclear correlation (HETCOR) spectra. The characteristic features of the mineral particle interfaces are the presence of non-apatitic hydrogen phosphate anions and water strongly bound to mineral crystal surfaces. 2D ^1^H-^31^P heteronuclear correlation (HETCOR) spectra resolve ^31^P environments according to the chemical shift of their spatially-close (<1 nm) protons (^1^H) and thus allow us to observe these key molecular components directly. Control samples exhibited the expected strong correlations with the maxima of the ^31^P peaks (δ(^31^P) ∼ 3ppm) correlating with proton chemical shifts of δ(^1^H) ∼ 0 ppm from hydroxyl groups in crystalline HAP, with ^1^H signals at δ(^1^H) ∼ 6.5 ppm, corresponding to ^31^Pat the surface of the HAP crystal lattice in close proximity to strongly-bound structural water at the mineral platelet surface and diffuse signal intensity between δ(^1^H) 8 - 17 ppm from hydrogen phosphate species in the amorphous hydrated platelet envelope as previously established for *in situ* bone mineral(*4*, *6*, *7*)(Fig. 2, C and D). OM samples exhibited the same spectral correlations indicating a similar mineral molecular structure despite the pathology in these samples (Fig. 2, C,D). HPP patient 1716 demonstrated a profound absence of structural water and amorphous phosphate species, with only an intense HAP mineral signature observed, with a narrow ^31^P linewidth compared to controls as in the 1D ^31^P spectra for these samples, indicating a mineral phase similar to microcrystalline HAP with very little hydrated interface (Fig. 2, C,D). HPP patient 0124 demonstrated a similar molecular structure by HETCOR NMR, but with an additional low intensity correlation at δ(^1^H) 4.6 ppm, a significantly lower ^1^H chemical shift than is observed for the strongly-bound platelet interfacial water in normal bone. This lower water ^1^H chemical shift indicates the presence of weakly bound water at the surface of mineral platelets in this patient’s bone mineral (Fig. 2, C,D).

Collectively, these NMR data indicate that the bone mineral molecular structure in HPP is distinct from that of healthy individuals and also from the mineral seen in other forms of OM. The highly crystalline, HAP-like mineral phase observed HPP patients is accompanied by a near-complete loss of structural water and amorphous hydrogen phosphate species that characterise the interfacial material between mineral platelets in normal bone mineral. Bone mineral is thought to form through sequential phase transformations(*44–46*). We hypothesized that the drastically different mineral structure in HPP bone reflects differential phase dynamics caused by the presence of high PPi concentrations in the bone mineralisation milieu, and therefore that excessive PPi impairs the normal regulation of calcium phosphate phase transitions during biomineralization, rather than simply inhibiting mineral nucleation as is currently widely believed.

### Inhibition of TNSALP does not alter early amorphous calcium phosphate mineral precursors during biomineralisation

To investigate the possibility of altered mineral phase dynamics in HPP, we first examined whether elevated PPi concentrations resulting from TNSALP inhibition could influence the earliest mineral phases that form during the biomineralisation process. Altering the structure or composition of the first-formed mineral phases could directly explain the differences in mature bone mineral molecular and nanostructure we observed in HPP bone and the altered microstructure of forming mineral on bone trabecular surfaces.

We induced matrix synthesis and mineralization in MC3T3-E1 osteoblasts using osteogenic medium over an extended time course while inhibiting TNSALP activity with the soluble inhibitor levamisole. In this model, the inorganic phosphate (Pi) required for mineral formation is generated through TNSALP-mediated hydrolysis of β-glycerophosphate present in the osteogenic medium. Inhibition of TNSALP therefore reduces that rate of Pi generation and is expected to slow the kinetics of mineral formation. If this were the only consequence of TNSALP inhibition, the mineral phase produced would remain unchanged. However, TNSALP inhibition also prevents the hydrolysis of PPi, leading to abnormally high PPi concentrations in the culture environment. These elevated PPi levels may, in turn, alter the phase of the mineral formed at any stage of the mineralization process. This experimental framework thus gives us an effective platform to assess whether PPi influences the earliest stages of bone mineral nucleation and transformation.

Mineralised nodules were generated in cultures upon stimulation with osteogenic media. Nodules were fewer in cultures treated with levamisole, but the cultures exhibited increased optical density at both 3 and 6 weeks of culture (Fig. 3, A). The molecular structure of the mineral particles generated with and without levamisole treatment was investigated using 2D ^1^H-^31^P ssNMR spectroscopy as for HPP patient bone samples. To ensure NMR signals are exclusively from the mineralized matrix and not cellular components, samples were decellularized prior to spectroscopic investigation. No change in collagen molecular structure was detected in ^13^C{^1^H} CP spectra of U-^13^C glycine- and L-proline-labelled samples (Supp. Fig. S3).

**Fig. 3.**
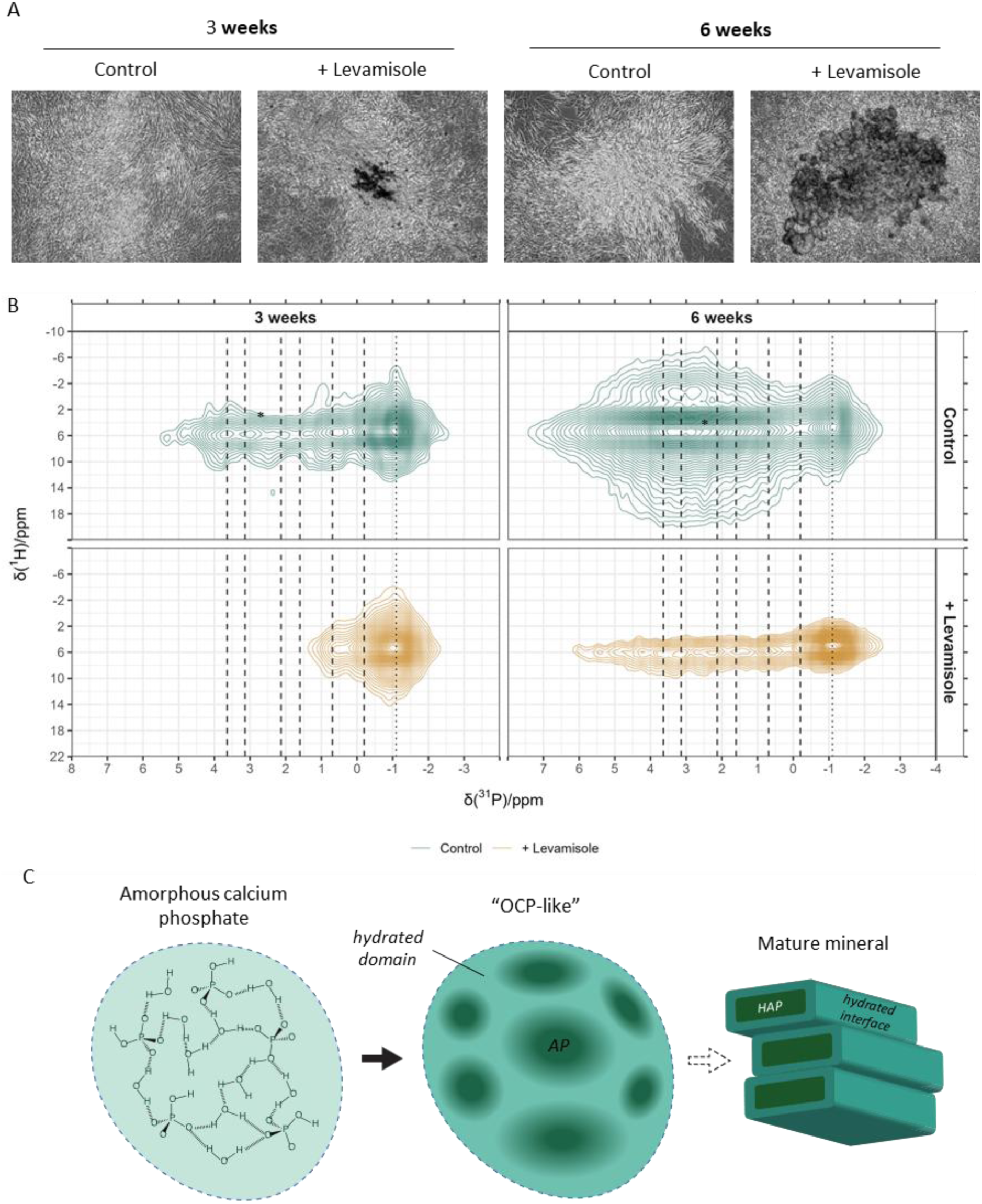
An in vitro model of hypophosphatasia (HPP) recapitulates altered calcium phosphate phase dynamics. **(A)** Phase contrast microscopy of MC3T3-E1 pre-osteoblasts cultured in osteogenic media ± levamisole for 3 or 6 weeks. **(B)** ^1^H-^31^P frequency-switched Lee-Goldburg heteronuclear correlation spectra of the same samples. Data are displayed with 40 contours between 15-100% normalised signal intensity. Dashed lines demonstrate expected chemical shifts of OCP-like ^31^P environments, while dotted lines demonstrate chemical shifts of the precursor ACP phase. **(C)** Schematic illustration of early calcium phosphate phase transformation during biomineralization, from an acidic ACP precursor, through a hydrated phase with OCP-like domains to mature bone mineral platelets.

The resulting 2D ^1^H-^31^P correlation spectra in control samples at 3 weeks of calcification show that the dominant signal in all cases is a broad ^31^P signal at δ(^31^P) -1.1 ppm correlated to a very broad ^1^H signal, centred at δ(^1^H) 5.0 - 5.6 ppm from strongly bound water, and extending to δ(^1^H) 8 - 11 ppm from ^1^H-^31^PO_4_^2-^ correlations, consistent with the presence of an amorphous mineral phase dominated by acidic HPO_4_^2-^ (Fig. 3, B; vertical dotted lines). We tentatively assign this signal to a precursor ACP mineral phase, widely believed to be the first step of bone mineral formation(*44–46*). The high intensity of this mineral component at the earliest stage of matrix mineralization shows that it is one of the first mineral phases to form; the broadness of the ^31^P signal strongly suggests it is part of an amorphous phase; and the correlation of this signal to a strongly-bound water signal is consistent with the expectation that ACP phases are highly hydrated. That this precursor phase in our *in vitro* model is dominated by acidic HPO_4_^2-^phosphatic sites is interesting as the earliest mineral formation event has been predicted to be formation of Posner or prenucleation clusters(*47*), loose networks of HPO_4_^2-^, Ca^2+^and water in the local mineralizing environment.

Other weaker correlation signals are from OCP-like ^31^P chemical environments: δ(^31^P) -0.2 ppm, 0.69 ppm, 1.6 ppm, 2.1 ppm, 3.1 ppm and 3.64 ppm correlating with a different strongly-bound water ^1^H signal at ∼ 6 ppm, confirming the OCP character of this intermediate mineral phase (Fig. 3, B; vertical dashed lines). By 6 weeks of culture these early mineral phases had largely been replaced with the expected signals from HAP-like ^31^P chemical environments: a broad peak at δ(^31^P) 2.8 - 3.3 ppm correlated with HAP hydroxyl proton environments at δ(^1^H) 0 ppm, 5.5 ppm (interfacial water) and 8 - 11ppm (interfacial hydrogen phosphate).

2D ^1^H-^31^P NMR spectra from levamisole-treated samples at 3 weeks calcification showed the same correlation signal from the ACP precursor mineral phase and a weak signal at δ(^31^P) 1.6 ppm correlated also correlated with water ^1^H (Fig. 3, B). At 6 weeks of culture, similar OCP-like signals are emerging to those seen for control cultures at the earlier 3-week time point (Fig. 3, B). These spectra for TNSALP-inhibited cultures are consistent with the expected slower mineral formation from slower generation of exogenous Pi, but more importantly show that the earliest mineral phases formed are highly similar to those in control samples, despite TNSALP inhibition presumably elevating PPi concentrations during mineral formation in these cultures.

These results support a model in which bone mineralisation involves early deposition of a highly hydrated, acidic ACP phase, which progressively transitions through phases containing OCP-like molecular structures before maturing into a more ordered bone mineral molecular architecture (Fig. 3, C). Importantly, inhibition of TNSALP does not appear to affect the form of these early mineral phases.

### Inorganic pyrophosphate alters the physiochemical dynamics of calcium phosphate mineral transformation

As inhibition of TNSALP does not appear to affect the early mineral phases that form on route to the final mature bone mineral, we hypothesised that it is elevated PPi present during mineral transformation that causes the formation of the high HAP crystallinity and disruption to interfacial hydrated layers between mineral crystals observed in HPP patient samples. To explore this with no confounding factors from cellular activity, we examined the effect of PPi on nanocrystalline HAP formation in solution. We used solid α-tricalcium phosphate (α-TCP) as the source of calcium and phosphate ions to ensure their constant concentration in solution, mimicking the *in vivo* calcification environment, and to avoid potentially confounding effects of other ions that could arise if we used separate calcium and phosphate salts as the reactants. Synthesis of HAP from α-TCP is also known to proceed via a transient OCP phase, replicating key aspects of calcium phosphate phase dynamics thought to occur during biomineral formation(*44–46*). Importantly, this synthetic route for nanocrystalline HAP is known to result in nanoscopic HAP layers interlayered with amorphous, hydrated non-apatitic layers mimicking the amorphous hydrated interfacial layers in *in vivo* bone mineral through dissolution-recrystallisation followed by hydrolysis(*12*, *48*, *49*). Thus, we synthesized HAP from α-TCP in the presence of increasing PPi concentrations to model the possible mineral products that could arise purely from chemistry in the absence of any directing influence from the organic components of the extracellular matrix (ECM) or other biomolecules.

TEM showed that control mineral samples demonstrated the expected thin platelet morphology of nanocrystalline HAP, mimicking *in vivo* bone mineral particle morphology with no preferred platelet orientation (Fig. 4, A). SAED exhibited characteristic reflections from the (002), (112) and (004) HAP lattice planes (Fig. 4, B). In contrast, mineral samples containing PPi presented as large flat plates in TEM and exhibited localised interference patterns at sites across the plates which increased in area with increasing PPi concentration (Fig. 4, A; white arrowheads).

**Fig. 4.**
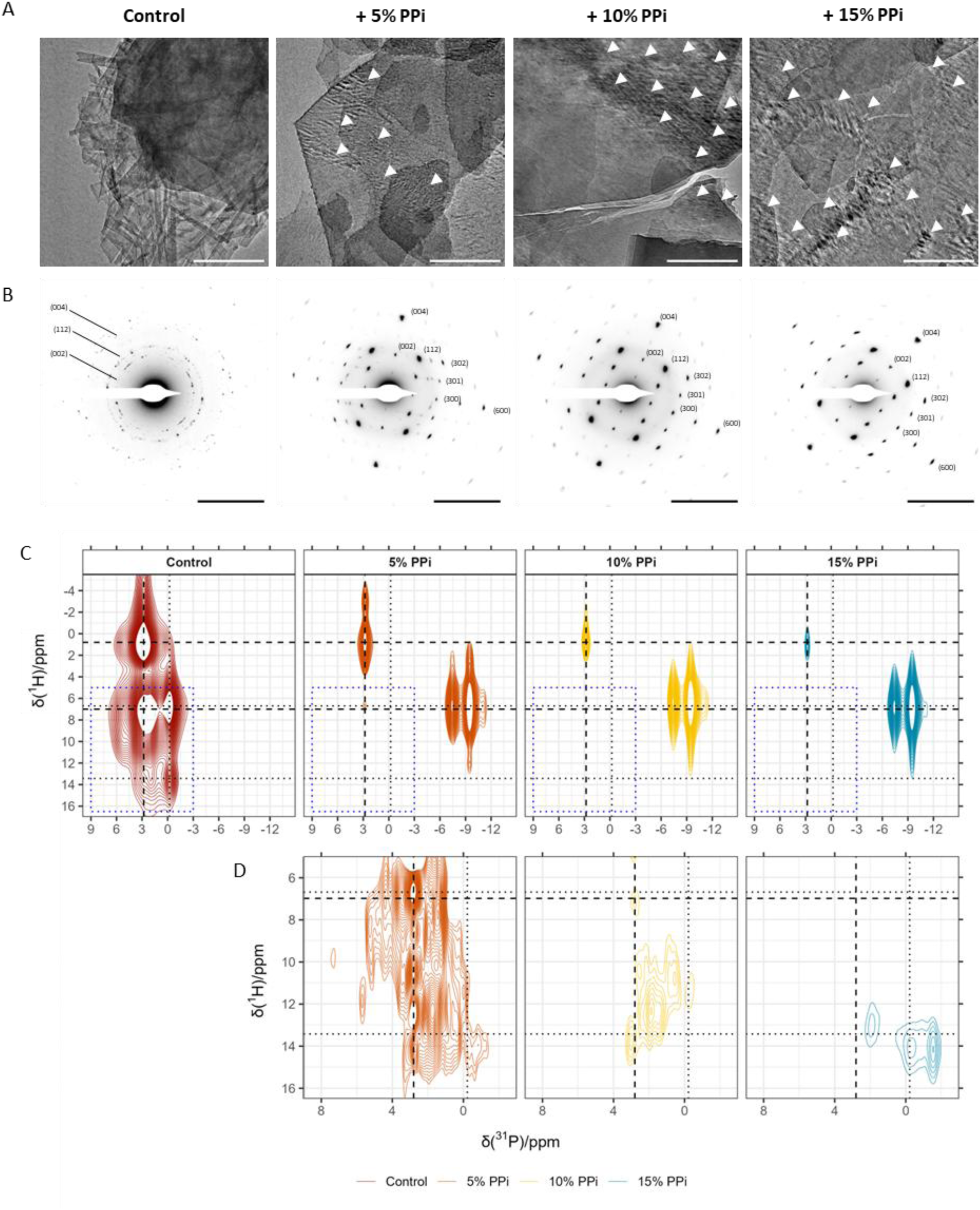
Characterisation of the nanostructure and molecular structure of synthetic bone mineral materials with incorporation of inorganic pyrophosphate (PPi). **(A)** Brightfield transmission electron microscopy. White arrowheads indicate interference patterns observed in the presence of PPi. Scale bars = xx nm. **(B)** Selected area electron diffraction (SAED) in the same samples as (A). Labels indicate hydroxyapatite Miller index of the corresponding reflection. Scale bars = 5 nm^-1^. **(C)** 2D ^31^P-^1^H frequency-switched Lee-Goldburg (FSLG) heteronuclear correlation (HETCOR) SSNMR spectra. Dashed lines indicate correlations characteristic for bone mineral and dotted lines characteristic for octacalcium phosphate (OCP). Spectra demonstrate a dominant HAP signal at δ(^1^H) ∼ 0 ppm, δ(^31^P) 2.8 ppm from OH- - PO_4_^2^-correlations; broad ^1^H signal δ(^1^H) 6 – 12 ppm correlated with the same δ(^31^P) 2.8 ppm signal; and residual OCP (δ(^31^P) -0.2 ppm; shoulder at 1.9 ppm correlated with water (δ(^1^H) ∼6 ppm) and HPO_4_^2^- (δ(^1^H) ∼13 ppm) ^1^H **(D)** Low intensity correlation signals in PPi-containing samples at chemical shifts within the blue dotted box in (D). FSLG spectra are plotted with (D) 40 contours between 5 – 40% normalised signal; (E) 40 contours between 0.6 – 5% normalised signal intensity.

SAED showed that these PPi-containing minerals consist of highly ordered HAP crystals, with patterns exhibiting characteristic HAP reflections which appeared to have a dual axis of symmetry through the diffraction pattern (Fig. 4, B, Supp. Fig. S4 and Supp. Table S2). Additional weak diffraction signals in +5% and +10% PPi samples were observed with d-spacings of 3.3 Å, 2.0 Å and 1.7 Å which could not be assigned, potentially indicating the presence of an additional crystalline phase at low PPi concentrations (Supp. Fig. S4 and Supp. Table S3). Darkfield TEM imaging further confirmed that interference patterns in these crystals represent regions where two twinned single crystals were sandwiched at a consistent orientation of 65°, generating a diffraction grating (Supp. Fig. S5). The presence of PPi during HAP nucleation therefore appears to induce long-range molecular order in HAP crystals and twins crystals at their {002} faces through an unknown mechanism.

We next examined the mineral molecular structure with SSNMR to determine the effect of PPi on the amorphous hydrated interlayers analogous to the HAP interfacial layers in *in vivo* bone mineral.2D ^1^H-^31^P HETCOR spectra (Fig 4, C,D) confirmed the mineral composition of control samples to be nanocrystalline HAP with associated amorphous, hydrated interlayers along with residual OCP (see figure legend for assignments), as expected for this HAP synthesis route. However, spectra from mineral samples synthesized in the presence of PPi are significantly different. First, the characteristic HAP signal (δ(^1^H) ∼ 0 ppm, δ(^31^P) 2.8 ppm from OH^-^ - PO_4_^3-^ correlations) is considerably narrower in the ^31^P dimension, suggesting significantly more long-range order of the orthophosphate anions in the HAP component of the mineral. Secondly, the correlation signals from amorphous, hydrated interlayers between HAP crystallites and from any OCP content (Fig. 4, D) are very weak even for the lowest PPi concentration (5 wt%) and substantially decrease in intensity with increasing PPi concentration. This suggests both a lack of amorphous hydrated interlayers around HAP crystallites and a phase transformation route to HAP that does not involve OCP, which indicates a highly altered mineral transformation route from the expected one.

The spectra also show clear additional signals from PPi ^31^P at δ(^31^P) -7.43 and -9.79 ppm (Fig. 4, C), demonstrating that PPi anions are contained within the solid precipitate. These ^31^P chemical shifts are consistent with the presence of a monoclinic calcium pyrophosphate tetrahydrate β (m-CPPTβ) hydrated pyrophosphate phase (*50*, *51*). These pyrophosphate ^31^P signals have a broad range of ^1^H correlations, including with water environments and hydrogen phosphate ^1^H, but not with HAP mineral hydroxyl groups (Fig. 4, C), indicating pyrophosphate anions are not close in space to these hydroxyl groups, but are close to strongly-bound water, consistent with an m-CPPTβ-like structure in the sample. Furthermore, 2D ^31^P-^31^P homonuclear radio-frequency-driven recoupling (RFDR) experiments in the +15% PPi sample also failed to demonstrate dipolar coupling between PPi and HAP ^31^P (Supp. Fig. S6). Interestingly however, the PPi and HAP orthophosphate ^31^P signals share a common small range of correlated ^1^H signals between δ(^1^H) 2 – 4 ppm, suggesting that the HAP and PPi-containing mineral components are structurally linked via these protons. Bulk power X-ray diffraction (pXRD) patterns from these samples also showed weak signals consistent with an m-CPPTβ component with a characteristic reflection at a d-spacing of 7.35 Å from the (100) plane(*51*), however these appeared broadened, suggesting a deformation or strain in the crystal lattice (Supp. Fig. S7). Taking these data together, we tentatively conclude that the m-CPPTβ component is bound to an HAP surface, causing strain in the m-CPPTβ lattice and potentially twinning HAP crystals at their {002} faces. Given the m-CPPTβ crystal structure includes layers of structural water at the crystal face, the HAP and PPi lattices may coalesce at this water layer.

These data support the radical alteration of calcium phosphate phase dynamics by PPi, shifting the system away from OCP-like transient intermediates and towards formation of a highly crystalline HAP, despite mineral formation seeming to still begin from an ACP phase. Thus, rather than PPi being an inhibitor of HAP formation, it changes the mechanism of HAP formation, leading to a substantially altered mineral composition, molecular and nano-structure, organization of mineral nanoparticles and spatial distribution of mineral. The striking absence of structural water and hydrogen phosphate species in the resulting mineral phase with increasing PPi concentration provides a compelling mechanistic link between elevated PPi in HPP and the absence of hydrated, amorphous mineral components observed in patient bone mineral and that this absence is due to the altered mechanism of HAP formation.

## Conclusions

Our findings demonstrate that extracellular PPi regulates the physiochemical dynamics of calcium phosphate phase transformations during biomineralization, slowing the kinetics of these transformations and resulting in highly ordered HAP crystals with a loss of hydrated interfacial layers. Previous work has detailed the composition of the highly hydrated and disordered interfacial phase between mineral platelets in bone(*4*, *6*, *7*). Here we show that this interface is a critical functional component of bone mineral molecular architecture and its loss as a result of elevated extracellular PPi in HPP patients severely compromises bone material function.

While the clinical symptoms associated with HPP have previously been characterised as an osteomalacic phenotype associated with hypomineralisation(*24*, *52*), here we show that HPP pathogenesis in severe compound heterozygous patients is a direct result of a defect in bone mineral chemistry and molecular architecture, distinct from other forms of OM. The pathophysiology of other rare bone conditions may therefore also lie in mineral molecular structure, or its molecular interface with collagen fibrils. Given the structural similarity between PPi and bisphosphonate drugs, the interference of pyrophosphate groups with mineral nucleation should also be considered as a potential root cause of fractures observed in patients treated with long-term bisphosphonates(*22*, *53*).

Finally, these findings suggest that controlled modulation of biomineralization could provide a powerful strategy for tuning mineral phase behaviour. Controlled alteration of PPi might allow deliberate manipulation of crystallinity and interfacial hydration, offering new approaches for engineering next generation biomimetic materials with tailored mechanical properties. More broadly, this work illustrates how biochemical control of local physiochemical kinetics can dictate the trajectory of non-equilibrium phase transformations in biological materials.

## Materials and methods

### Materials

LR-White resin, α-Tricalcium phosphate (α-TCP; CAS: 7758-87-4), sodium hydroxide, sodium pyrophosphate dibasic (CAS: 7758-16-9), L-ascorbic acid (CAS: 50-81-7), β-glycerophosphate (βGP, CAS: 154804-51-0), dexamethasone (CAS: 50-02-2), levamisole hydrochloride (CAS: 16595-80-5), Triton X-100, ammonium hydroxide (NH_4_OH), Tris-HCl, magnesium chloride and calcium chloride were purchased from Sigma-Aldrich.

### Human clinical investigations, sampling and histomorphometry

Patients were recruited from the Eastern Region Rare Bone clinic, (ErBON), part of the NHS Rare Diseases Collaborative Network (RDCN) at Addenbrooke’s hospital, Cambridge, UK. Patients were invited to participate in the research study, ’Characterisation of the mineral content of bone in hypophosphatasia" (Integrated Research Application System ID 304767), Cambridge NIHR Biomedical Research Centre (BRC study code RG64245). All ErBON patients with a clinical diagnosis of hypophosphatasia are entered into the COBRA database (Cambridge Outpatients Bone Registry), have blood taken for whole genome sequencing (WGS), dual X-ray absorptiometry (DXA) of the hip and spine (Hologic) and GE Lunar Quantitative Computed Tomography using a bone-equivalent phantom (Mindways QCT Pro) for measurement of spinal trabecular bone density. Participants had extensive serum bone profile blood tests including alkaline phosphatase.

Patients underwent a standard diagnostic 7.5 mm diameter Bordier transiliac bone biopsy from a site 2 cm inferior and posterior to the anterior-superior iliac spine, according to a previously described technique(*54*). One Jamshidi needle core for routine decalcified examination was also taken, in order to exclude malignancy. Specimens were fixed in 70% ethanol, dehydrated through an ethanol series, and embedded in LR-White resin. 10 um undecalcified sections were obtained using a microtome with a tungsten carbide knife. Undecalcified sections were stained with either von Kossa or toluidine blue stains while decalcified sections were stained with hematoxylin and eosin according to standard protocols(*55*). Bone histomorphometric analysis was conducted in trabecular bone using the semiautomatic method provided by Bioquant Osteo II software (Bioquant image analysis corporation). All histomorphometric parameters were reported using the nomenclature recommended by the American Society for Bone and Mineral Research(*56*). Histomorphometry data were compared to standard reference data to calculate Z-scores(*57*). The bone biopsies obtained from the two compound heterozygous patients (patients 0127 and 1716) with hypophosphatasia were compared to those from three patients with other forms of osteomalacia (one with tumor-induced osteomalacia and two with X-linked hypophosphatemia; patients 0120, 0718 and 1816), as well as to three reference patients who were found to have normal mineralization (patients 0517, 0817 and 1517). All samples have generic research consent.

### Synthesis of model bone mineral

To prepare a control material α-TCP (0.8219 g, 2.65 mmol) was added to 50 ml distilled water and the pH adjusted to 6.5 by dropwise addition of 0.5 M NaOH solution. The mixture was incubated with stirring (120 RPM) at 37°C for 10 days. The resultant precipitate was filtered under gravity and air-dried overnight. The product was obtained as a white solid (0.798 g).

To prepare mineral in the presence of inorganic pyrophosphate (PPi), the same method was used. Following addition of α-TCP 5% (0.029 g, 0.1325 mmol), 10% (0.0588 g, 0.2650 mmol) or 15% (0.0882 g, 0.3975 mmol) molar equivalent sodium pyrophosphate dibasic (Na_2_H_2_O_2_O_7_) was added to 50 ml distilled water and the pH adjusted to 6.5. Following incubation and filtration the products were obtained as white solids (0.743 g, 0.770 g and 0.707 g respectively). Minerals were stored at room temperature prior to analysis.

### Tissue culture and generation of in vitro extracellular matrices

Reagents for tissue culture were purchased from Gibco unless otherwise indicated.

The mouse pre-osteoblast MC3T3-E1 cell line (ATCC) was used to generate mineralised collagen I extracellular matrix (ECM). Cells were cultured at 37°C in a humidified atmosphere with 5% CO_2_ in complete growth media comprising α-Minimum Essential Media (αMEM) with nucleosides supplemented with 10% foetal bovine serum (FBS; PAN Biotech), 5 units/ml penicillin, 1 μg/ml streptomycin and 0.292 mg/ml L-glutamine. FBS was heat-inactivated prior to addition to growth media by incubation at 60°C for 2h. Complete growth media was refreshed every 2-3 days.

To induce extracellular matrix production and mineralisation, growth media was supplemented at cell confluence with 50 μg/ml L-ascorbic acid, 5 mM βGP and 10 nM dexamethasone, and cells cultured for 21 or 42 days, with osteogenic media refreshed every 2-3 days. For inhibition of tissue non-specific alkaline phosphatase (TNSALP) *in vitro*, osteogenic media was supplemented with 100 μM levamisole. For ^13^C isotopic enrichment of *in vitro* ECM, osteogenic media was further supplemented with 50 μg/ml U-^13^C glycine and 40 μg/ml U-^13^C L-proline (Cambridge Isotope Laboratories) for the entire post-confluence culture period. Phase contrast images during culture were acquired using a Evos XL Core (Thermo Fisher Scientific) microscope.

### Decellularisation of in vitro extracellular matrices

To remove cells from mineralised *in vitro* mineralised ECMs we used a detergent decellularization method. Mineralised matrices were generated as above for either 21 or 42 days. Osteogenic media was removed and cells washed in Dulbecco’s phosphate buffered saline (DPBS; Gibco). Matrices were frozen in DPBS and maintained at -20°C overnight. Samples were allowed to thaw for 30min at room temperature, washed in DPBS, then DBPS was removed and treated with 0.5% Triton X-100; 20 mM NH_4_OH in DPBS for 10 min at 37°C. Samples were decanted to a 50 ml Falcon tube, an equal volume of DPBS added, and incubated overnight at 4°C. Detergent buffer was discarded and samples washed thoroughly in DPBS. Samples were digested in 10 μg/ml DNAse I (Roche) in DNAse working buffer (100 mM Tris-HCl; 25 mM MgCl_2_; 1 mM CaCl_2_, pH 7.6) at 37°C for 30 min. Matrices were washed well in DPBS before preparation for downstream analysis.

### Scanning electron microscopy

Bone biopsies were prepared by polishing the surface of the resin block with sequential grade diamond polishing strips (Agar Scientific). Blocks were mounted onto aluminium sample stubs with adhesive carbon tabs (Agar Scientific) and electrical connection established with the sample surface using copper tape (Electron Microscopy Sciences). The sample surface was coated with 10 nm carbon using a Q150T ES carbon coater (Quorum Technologies).

Samples were imaged with a CLARA 2 (TESCAN) field emission gun scanning electron microscope (SEM) equipped with a low-energy annular backscattered electron (BSE) detector and operating at 10 kV and a working distance of 10mm. Energy-dispersive X-Ray spectroscopy (EDS) spectra were collected using an X-maxN 80 detector (Oxford Instruments) at the same conditions.

Images were processed using Fiji(*58*). Statistical analysis of EDS data was performed in R using an analysis of variance (ANOVA) followed by post-hoc testing using pairwise Tukey honest significant difference (HSD) tests.

### (Scanning) transmission electron microscopy

For transmission electron microscopy (TEM) and scanning-transmission electron microscopy (STEM) in human bone biopsies, ultrathin (90 nm) sections were prepared from biopsies using an EM UC7 ultramicrotome (Leica) equipped with a 45° Ultra diamond knife (Diatome) and mounted onto 200 mesh carbon and formvar-coated copper grids (Agar Scientific). To prepare model bone minerals, a small amount of material was resuspended in 200 μl absolute ethanol and sonicated for 30 s. The resulting suspension was pipetted (10 μl) onto grids as above and incubated at room temperature for 2 min, before excess ethanol was removed with filter paper and grids briefly air dried.

Microscopy was performed using a Talos F200X G2 FEG microscope (Themo Fisher Scientific) equipped with a Ceta-M CMOS camera (Thermo Fisher Scientific) and operating at 200 kV. Selected area electron diffraction (SAED) images were acquired using a 40μm selected area aperture and at a camera length of 1.3 m. Darkfield TEM was performed by first acquiring SAED patterns as above. Single diffraction signals were isolated using a 10 μm objective aperture and darkfield images acquired with the selected area aperture retracted. STEM images were recorded using a high-angle annular darkfield (HAADF) detector with a pixel dwell time of 10 μs.

SAED images were indexed by drawing rings where 2 equivalent diffraction spots across the central spot could be identified and calculating plane d-spacing in real space. Spacings were compared to hydroxyapatite (HAP) reference values(*59*). In single crystal diffraction patterns plane identification by d-spacing was verified by measuring the angle between at least 2 confirmed planes, calculated using the following equation for hexagonal crystals, where any 2 planes are defined by their Miller indices (h*_1_k_1_l_1_*) and (*h2k_2_l_2_*):

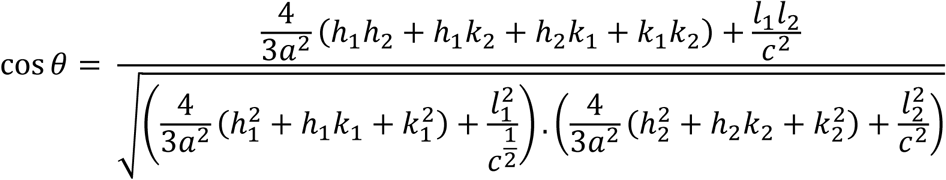

Where 𝑎 and 𝑐 are the crystal unit cell parameters, assumed to be 𝑎 = 𝑏 = 9.432 Å and 𝑐 = 6.881 Å in HAP.

Images were processed using Fiji(*58*).

### Solid state nuclear magnetic resonance spectroscopy

Consumables for solid state nuclear magnetic resonance (SSNMR) spectroscopy were purchased from Bruker.

For SSNMR experiments in human bone biopsies, 10 μM serial sections of biopsies were trimmed of excess resin and packed into 4 mm Kel-F inserts within 4mm ZrO_2_ rotors and capped with Kel-F drive caps. For synthetic model bone mineral, samples were packed directly into 4 mm rotors. For *in vitro* mineralised ECM, samples were centrifuged at 300 x g for 3 min at room temperature and bulk water removed. Hydrated matrices were packed into 3.2 mm rotors and capped with a VESPEL drive cap. Rotors were snap frozen on dry ice and stored at -20°C prior to analysis.

In bone biopsy and material samples, SSNMR was performed using an AVANCE Neo 400MHz (9.4T) ^1^H Larmor frequency wide bore spectrometer (Bruker) equipped with a 4mm HXY magic angle spinning (MAS) probe (Bruker) operating at 10kHz MAS at room temperature. In ECM samples, SSNMR was performed using an AVANCE Neo 600 MHz (14.1T) ^1^H Larmor frequency standard bore spectrometer (Bruker) equipped with a 3.2 mm HCP Efree MAS probe (Bruker) operating at 12 kHz MAS at a corrected sample temperature of -10°C. Sample temperature under MAS was corrected using the temperature-dependent T_1_ relaxation of KBr at the same conditions. ^31^P chemical shifts were referenced to crystalline HAP at 2.8 ppm (relative to 85% phosphoric acid at 0 ppm), ^1^H shifts in the same sample to 0 ppm (relative to tetramethylsilane (TMS) at 0 ppm), and ^13^C shifts to glycine C_α_ at 43.1 ppm (relative to TMS at 0 ppm).

In all experiments ^1^H 90° pulses were performed at 100 kHz and data were acquired with SPINAL64 ^1^H decoupling at 100 kHz. The recycle delay for ^1^H pulse-acquire 1D spectra, 1D ^31^P/^13^C{^1^H) cross polarisation (CP) and 2D ^1^H-^31^P heteronuclear correlation (HETCOR) spectra was 2.5 s. CP spectra were acquired with a ramped 50-100 or 70-100 ^1^H contact pulse and a square X pulse at contact times ranging from 200 μs to 10 ms depending on the experiment. The Frequency-Switched Lee-Goldburg (FSLG) pulse sequence was used to record ^1^H-^31^P HETCOR spectra with Lee-Goldburg homonuclear ^1^H decoupling during t_1_ evolution(*60*). All FSLG experiments were performed with a contact time of 2 ms. In bone biopsies, FSLG spectra were acquired with 1024 scans at 32 t_1_ increments; 256 scans at 64 t_1_ increments in model bone minerals; and 2048 scans at 32 t_1_ increments in *in vitro* ECM samples. To correct chemical shift scaling in the indirect dimension of FSLG spectra, ^1^H-^1^H homonuclear correlation spectra were recorded with identical FSLG offsets and delays and the increment for delay adjusted to give a perfect diagonal with the direct dimension, as previously described(*61*). In ^31^P direct polarisation spectra, the recycle delay was 10 s, and 90° pulses were performed at 80 kHz (mineral materials) or 86 kHz (ECM samples). In ^31^P-^31^P homonuclear radio-frequency-driven recoupling (RFDR) experiments, an initial ^31^P{^1^H} CP magnetization transfer was performed with a contact time of 10 ms as above and RFDR performed with recoupling times between 20 – 200 ms. ^31^P 90° and 180° pulses were performed at 72 kHz and spectra were recorded with 256 scans at 256 t_1_ increments.

In bone biopsies, 1D ^31^P CP spectral peaks were fitted with a Lorentzian lineshape using a nonlinear least squares approach according to the function:

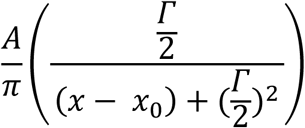

Where 𝐴 is peak amplitude, 𝛤 is peak full-width at half maximum, 𝑥 is chemical shift and 𝑥_0_ is the peak isotropic chemical shift.

NMR data processing was performed in R.

### Powder X-ray Diffraction

Powder X-ray diffraction (XRD) data were collected on a Malvern Panalytical Empyrean instrument, equipped with an X’celerator Scientific detector using non-monochromated CuKα radiation (λ = 1.5418 Å). The sample was placed on a glass sample holder and measured in reflection geometry with sample spinning. The data were collected at room temperature over a 2θ range of 10 to 70°, with an effective step size of 0.02° and a total collection time of 1 hour. XRD data were aligned and background corrected using the powdR package(*62*) for R and expressed as interplanar d-spacing.

## Acknowledgements

The authors gratefully acknowledge Mr Phillip Johnston, Mr Stephen McDonnell and Dr Gavin Clunie who assisted to diagnosis, clinical care and sampling of patients. The authors also acknowledge valuable technical assistance from Dr Heather Greer and Dr Christopher Truscott. S.D. and M.J.D. were supported by a European Research Council Advanced Grant (101019499). Instrumentation used in this work was supported by the Engineering and Physical Sciences

Research Council [EP/P030467/1].

## List of Supplementary Materials

1. Supplementary Figures S1-S7
2. Supplementary Tables S1-S3

## Supplementary materials

### Supplementary figures

**Supplementary Fig. S1.**
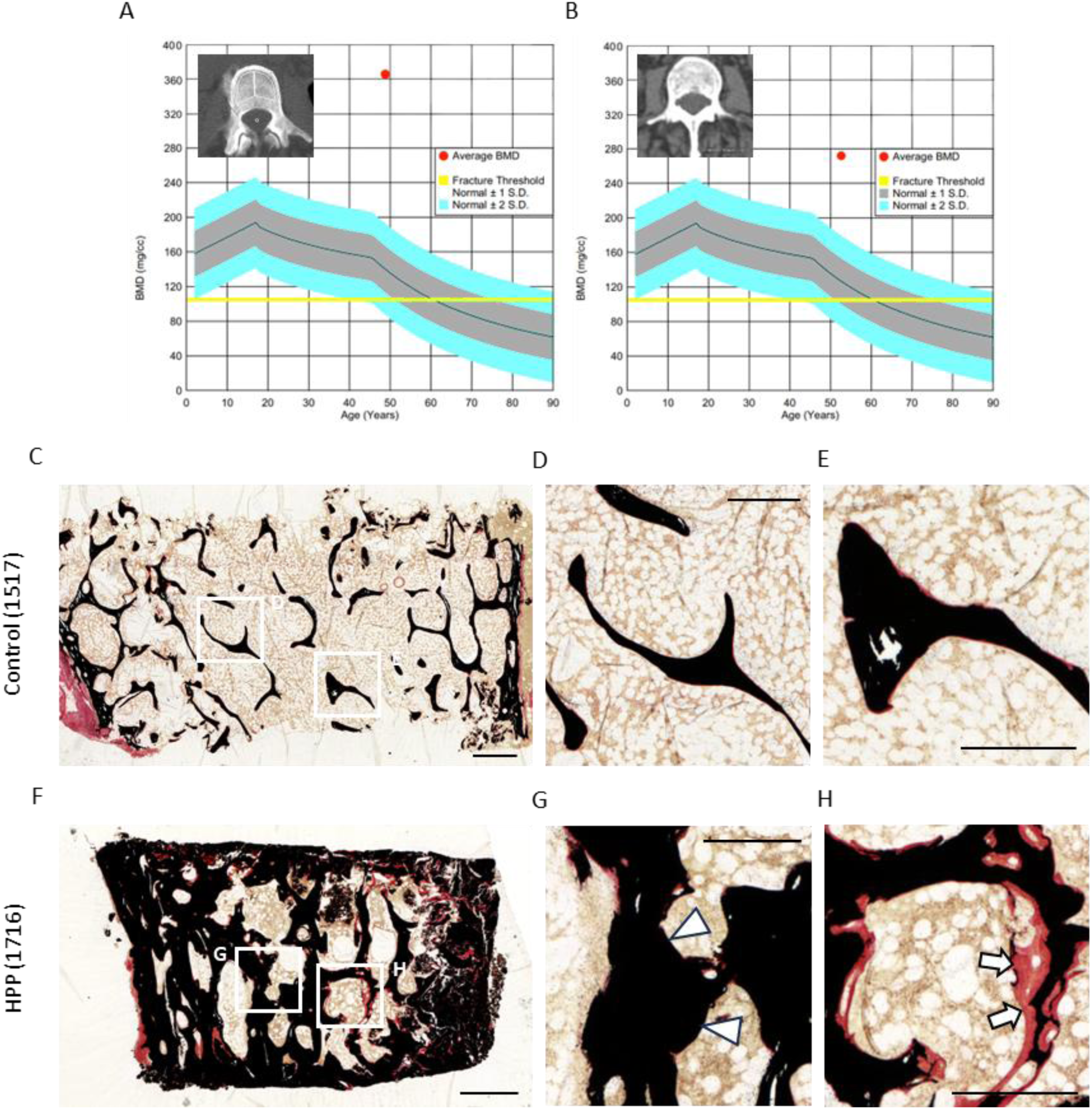
Clinical investigations in human patients diagnosed with hypophosphatsia (HPP) and other forms of osteomalacia. (A,B) Quantitative computed tomography (qCT) in patient 0124 (A) and patient 1716 (B) at L3 demonstrating a volumetric bone mineral density (vBMD) of 374mg/cm^3^ and 272mg/cm^3^ respectively. (C-H) Undecalcified transiliac crest bone biopsy histopathology with Von Kossa staining. HPP samples demonstrate simultaneous trabecular sclerosis (white arrowheads) and hyperosteoidosis (white arrows). Scale bars = 1000µm (C,F); 500µm (D-E,G-H).

**Supplementary Fig. S2.**
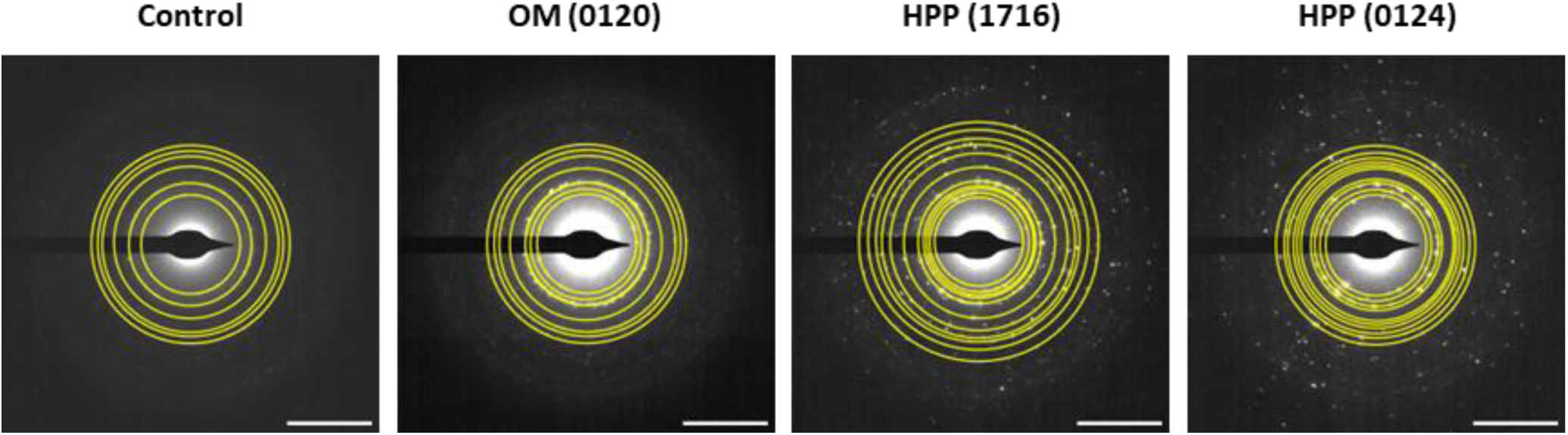
Full diffraction indexing of selected area electron diffraction (SAED) images in human patient bone biopsies. Patterns were indexed by drawing rings where 2 equivalent diffraction signals could be observed across the central spot, the interplanar d-spacing calculated and comparison with hydroxyapatite reference values(60).

**Supplementary Fig. S3.**
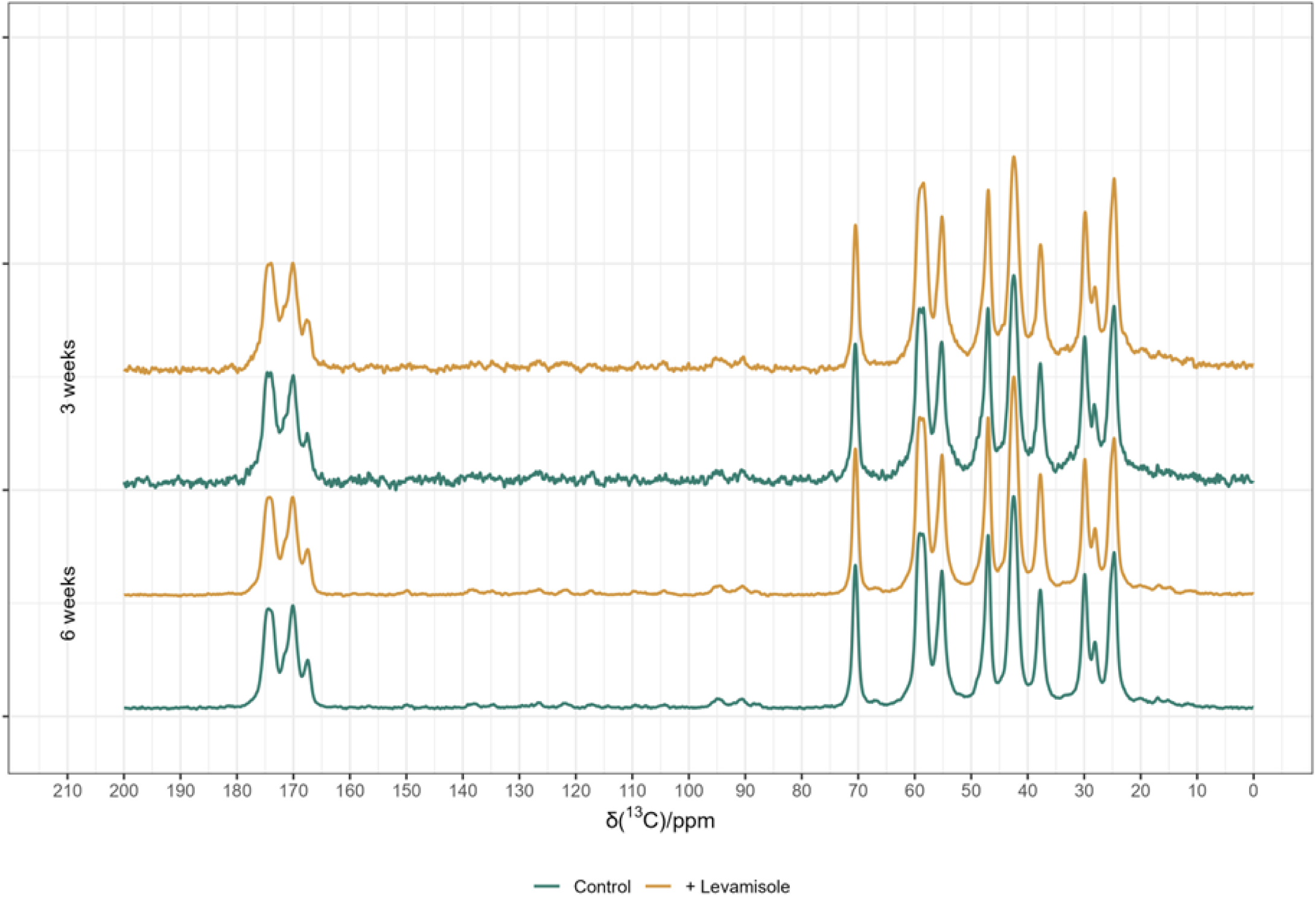
^13^C{^1^H} solid state nuclear magnetic resonance spectroscopy (SSNMR) spectra of in vitro decellularised mineralised matrices. Collagen signals from the mineralised matrix enriched by bioincorporation of U-^13^C labelled glycine and L-proline during culture. Signal intensity normalised to the Hyp C_γ_signal at δ(^13^C) 70.5 ppm.

**Supplementary Fig. S4.**
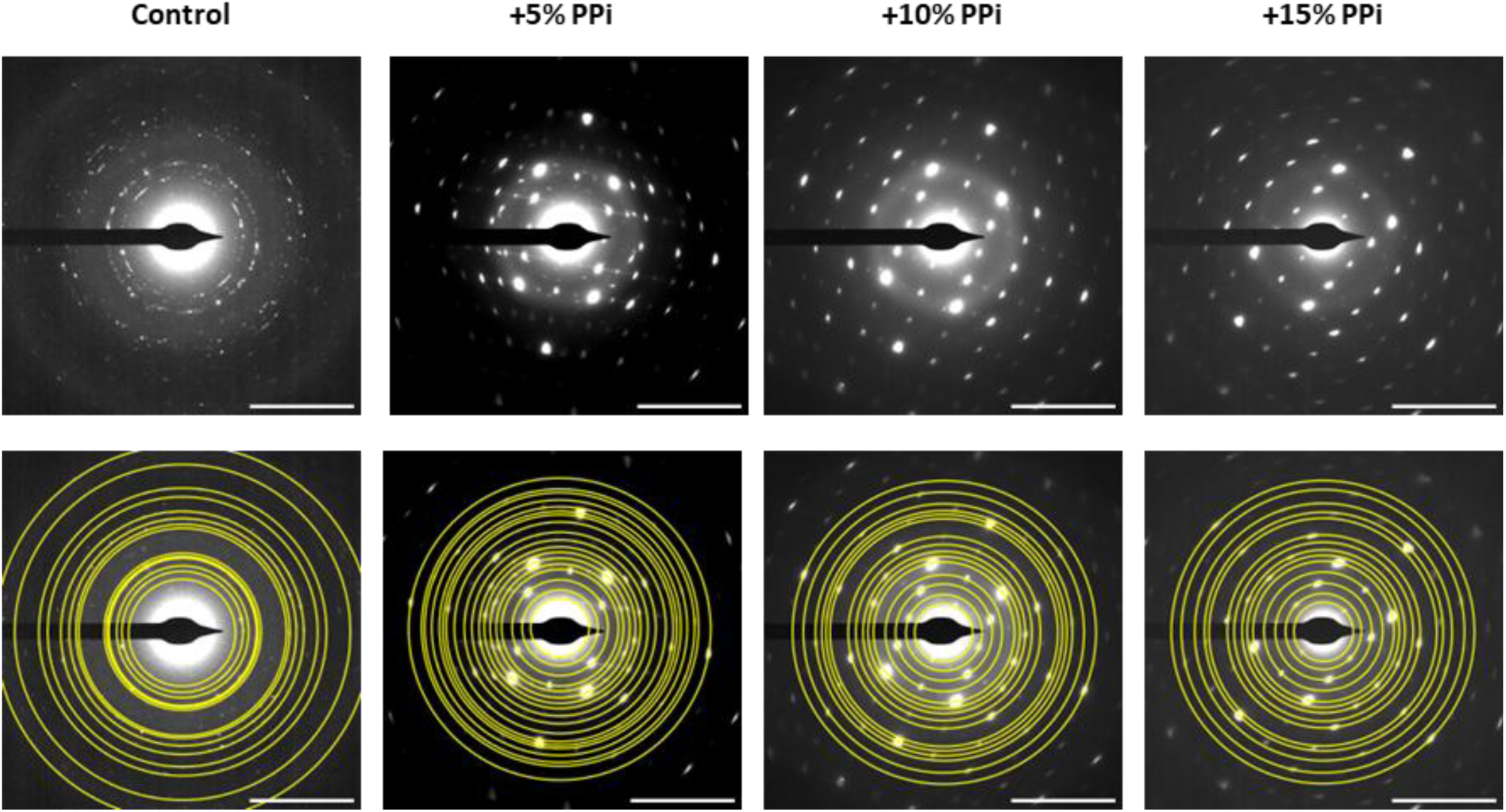
Full diffraction indexing of selected area electron diffraction (SAED) patterns in synthetic model bone mineral samples. Patterns were indexed by drawing rings where 2 equivalent diffraction signals could be observed across the central spot, the interplanar d-spacing calculated and comparison with hydroxyapatite reference values(60). Assigned planes were verified by confirming the observed angle between that signal and a previously confirmed signal (see Materials and Methods).

**Supplementary Fig. S5.**
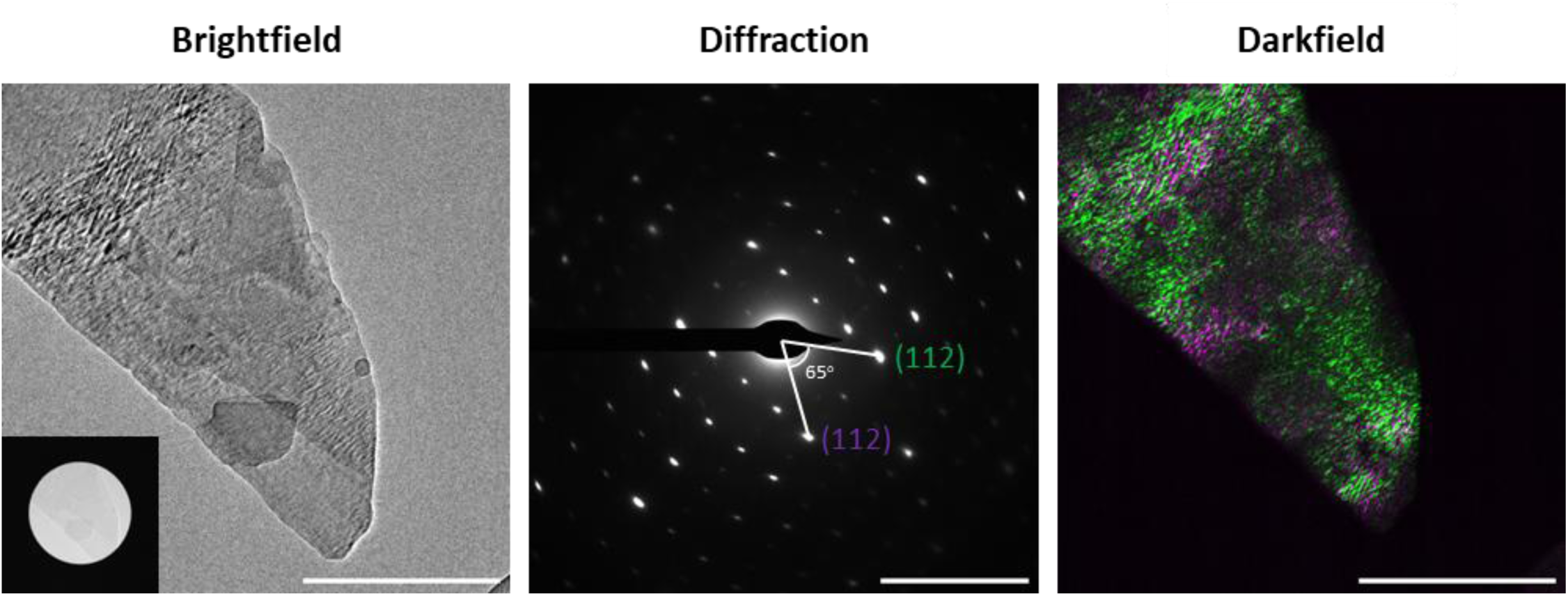
Darkfield transmission electron microscopy of synthetic model bone mineral samples. Imaging performed in a +15% inorganic pyrophosphate (PPi) sample. Selected area electron diffraction (SAED) performed with a 40 μm selected area aperture (brightfield; inset). Darkfield performed with a 10 μm objective aperture at labelled (112) reflections. Darkfield images are coloured according to isolated diffraction reflection. Scale bars = 500nm (brightfield and darkfield); 5nm^-1^ (diffraction).

**Supplementary Fig. S6.**
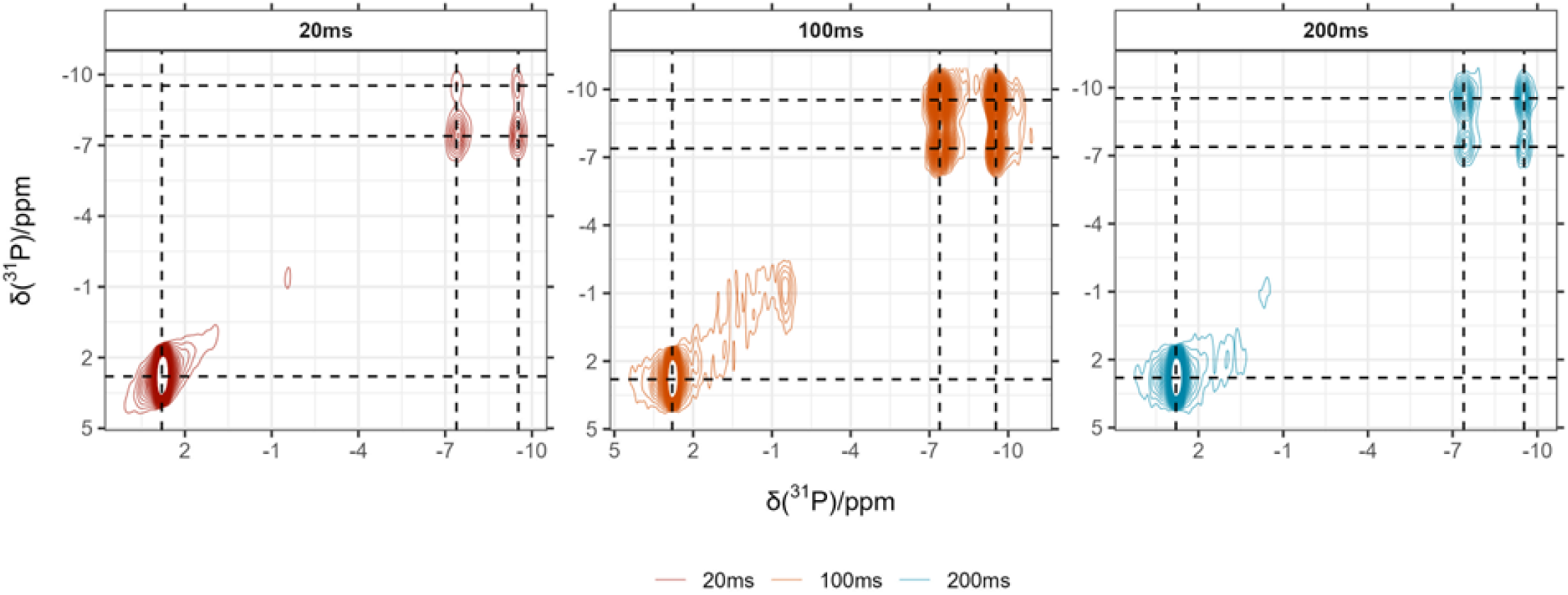
^31^P-^31^P homonuclear cross polarisation ^31^P{^1^H} radio-frequency-driven recoupling (RFDR) spectra in a + 15% pyrophosphate sample. Spectra demonstrate dipolar coupling (and therefore spatial proximity) between pyrophosphate ^31^P at δ(^31^P) -7.4 and -9.5 ppm, but not between pyrophosphate and hydroxyapatite orthophosphate ^31^P (dashed lines). Spectra performed with a cross polarisation contact time of 10 ms and recoupling times between 20 – 200 ms. Spectra are plotted with 64 contours between 1 – 40% normalised signal intensity.

**Supplementary Fig. S7.**
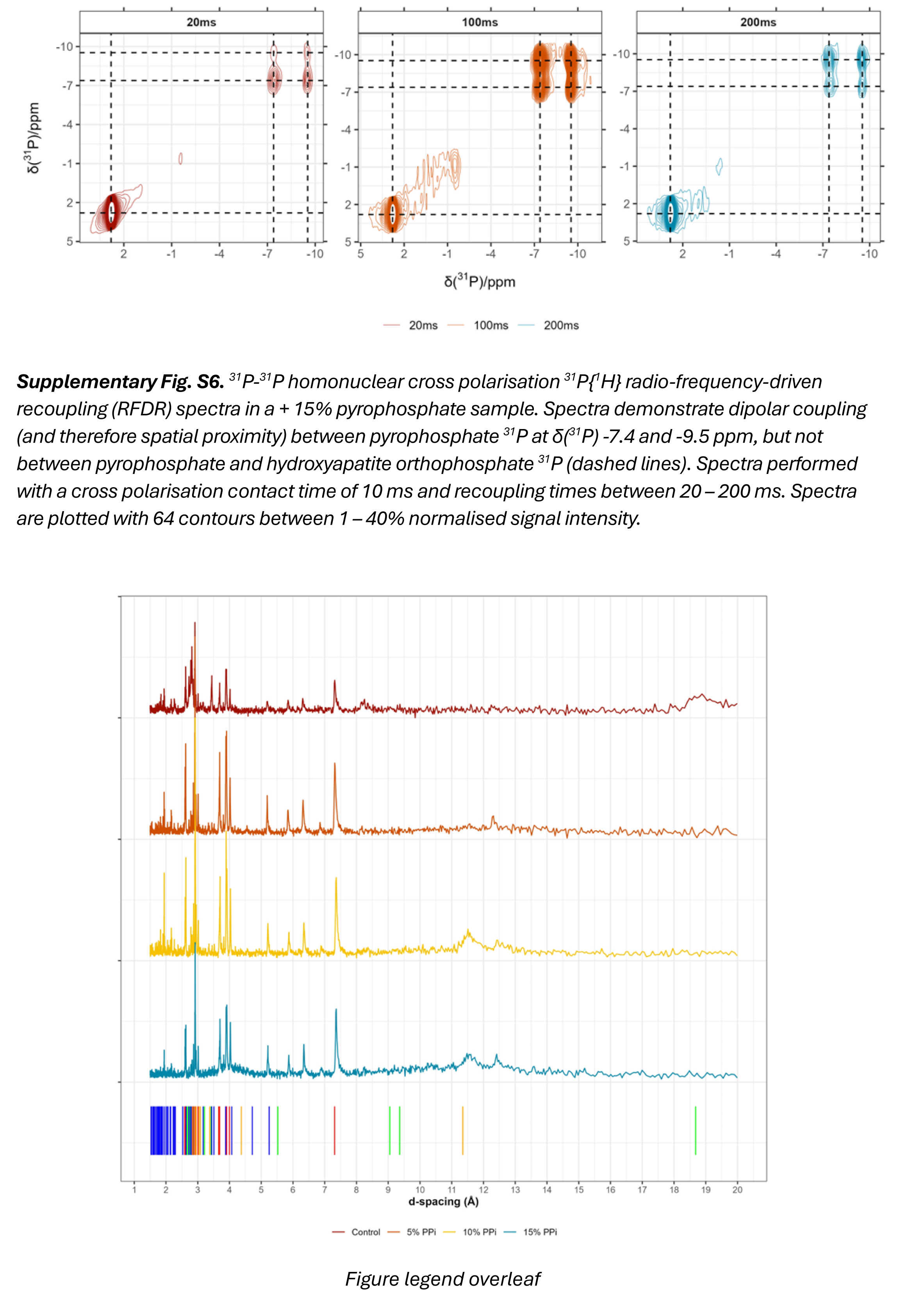
Powder X-ray diffraction (pXRD) of model bone mineral samples. Data are aligned and background corrected and expressed as interplanar d-spacing. Reference d-spacings for planes in α-tricalcium phosphate (αTCP; red), octacalcium phosphate (OCP; green), hydroxyapatite (HAP; blue) and monoclinic calcium pyrophosphate tetrahydrate β (m-CPPTβ; orange) are indicated below spectra (51,60).

### Supplementary tables

**Supplementary Table S1:** Patient undecalcified trans-iliac crest bone biopsy histomorphometry. Z-scores are calculated according to reference standard data(*56*, *57*).

| Patient | Diagnosis | BV/TV (%) | BV/TV Z score | OV/BV (%) | OV/BV Z score | OS/BS (%) | OS/BS Z score | MO.Th (μm) | MOTh Z score | Tb.Th (μm) | Tb.Th Z score | Tb.N (count/mm) | Tb.N Z score | Tb.Sp (μm) | Tb.Sp Z score |
| --- | --- | --- | --- | --- | --- | --- | --- | --- | --- | --- | --- | --- | --- | --- | --- |
| 0517 | Control | 23.2 | 0.25 | 0.1 | -2.5 | 1 | -2.87 | 3.67 | -1.47 | 130 | -0.24 | 1.78 | 0.21 | 431 | -0.8 |
| 0817 | Control | 21.6 | 0.71 | 0.7 | -1.55 | 4.3 | -2.68 | 11.18 | 0.65 | 133 | -0.19 | 1.63 | 0.33 | 484 | -0.54 |
| 1517 | Control | 23.45 | 0.29 | 0.3 | -2.33 | 2.65 | -2.56 | 5.94 | -0.89 | 71.96 | -2.56 | 1.71 | 0.03 | 448 | -0.64 |
| 1816 | XLH | 45.25 | 4.09 | 17 | 14.1 | 95 | 19.66 | 17.6 | 2.24 | 207 | 2.9 | 2.21 | 1.27 | 248 | -1.96 |
| 0718 | XLH | 43.05 | 3.73 | 26 | 23.1 | 96.5 | 20.02 | 23.46 | 3.67 | 167 | 1.16 | 2.59 | 2.23 | 221 | -2.15 |
| 0120 | TIO | 26.5 | 1.06 | 30.5 | 27.6 | 63 | 11.85 | 33.82 | 6.2 | 137 | -0.15 | 1.96 | 0.64 | 378 | -1.06 |
| 1716 | HPP | 44.8 | 4.71 | 9.15 | 6.14 | 45.45 | 10.6 | 18.33 | 2.81 | 157 | 0.43 | 2.88 | 3.45 | 192 | -2.7 |
| 0124 | HPP | 41.4 | 4.12 | 44.6 | 37.22 | 78.3 | 21.19 | 31.59 | 6.48 | 100.75 | -0.98 | 4.11 | 6.52 | 142.84 | -3.07 |

**Supplementary Table S2:**
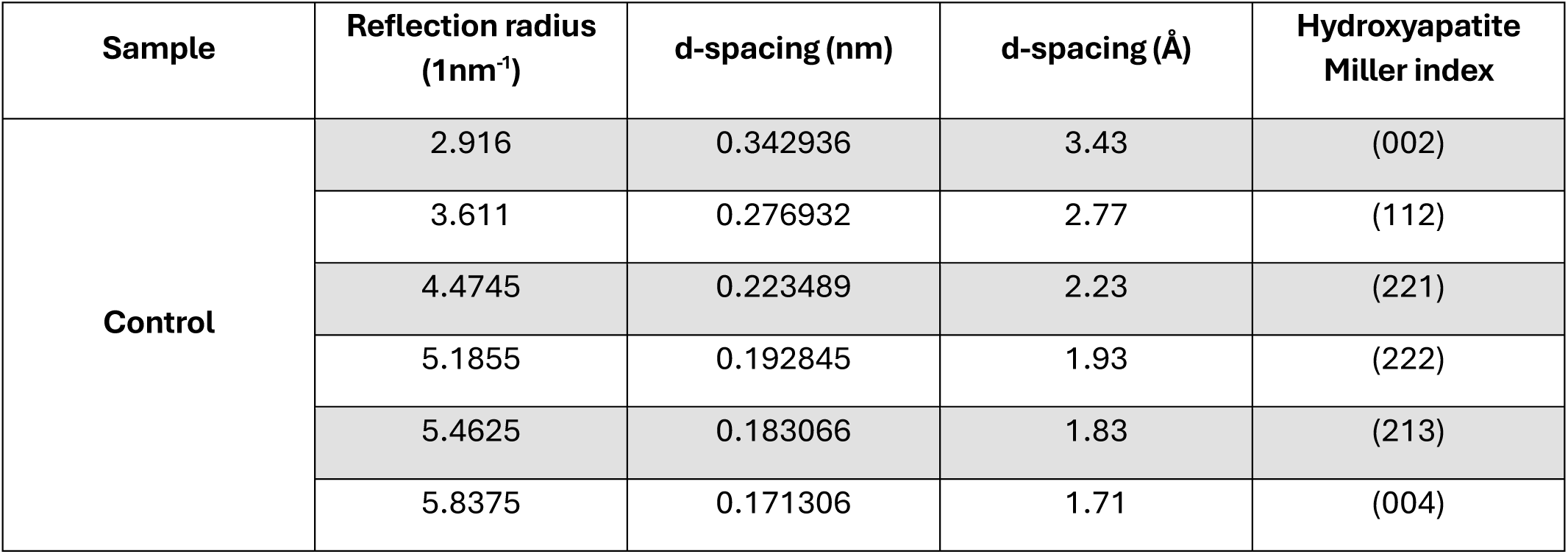
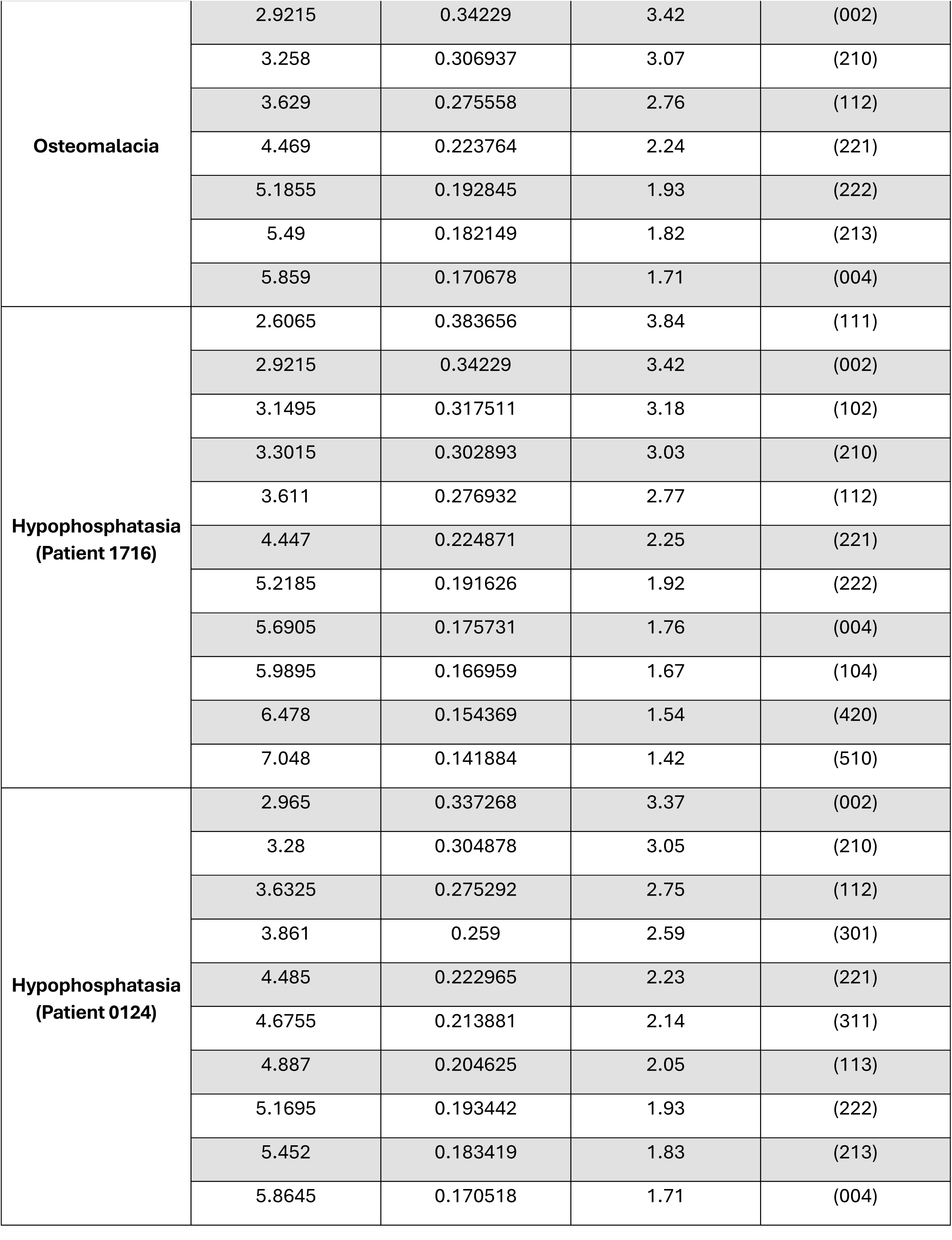
Full diffraction indexing of selected area electron diffraction (SAED) patterns in human trans-iliac crest bone biopsies. Miller indices are assigned according to reference standard X-ray data in hydroxyapatite (*59*).

| Sample | Reflection radius (1nm<sup>-1</sup>) | d-spacing (nm) | d-spacing (Å) | Hydroxyapatite Miller index |
| --- | --- | --- | --- | --- |
| Control | 2.916 | 0.342936 | 3.43 | (002) |
|  | 3.611 | 0.276932 | 2.77 | (112) |
|  | 4.4745 | 0.223489 | 2.23 | (221) |
|  | 5.1855 | 0.192845 | 1.93 | (222) |
|  | 5.4625 | 0.183066 | 1.83 | (213) |
|  | 5.8375 | 0.171306 | 1.71 | (004) |
| **Osteomalacia** | 2.9215 | 0.34229 | 3.42 | (002) |
|  | 3.258 | 0.306937 | 3.07 | (210) |
|  | 3.629 | 0.275558 | 2.76 | (112) |
|  | 4.469 | 0.223764 | 2.24 | (221) |
|  | 5.1855 | 0.192845 | 1.93 | (222) |
|  | 5.49 | 0.182149 | 1.82 | (213) |
|  | 5.859 | 0.170678 | 1.71 | (004) |
| **Hypophosphatasia**<br>**(Patient 1716)** | 2.6065 | 0.383656 | 3.84 | (111) |
|  | 2.9215 | 0.34229 | 3.42 | (002) |
|  | 3.1495 | 0.317511 | 3.18 | (102) |
|  | 3.3015 | 0.302893 | 3.03 | (210) |
|  | 3.611 | 0.276932 | 2.77 | (112) |
|  | 4.447 | 0.224871 | 2.25 | (221) |
|  | 5.2185 | 0.191626 | 1.92 | (222) |
|  | 5.6905 | 0.175731 | 1.76 | (004) |
|  | 5.9895 | 0.166959 | 1.67 | (104) |
|  | 6.478 | 0.154369 | 1.54 | (420) |
|  | 7.048 | 0.141884 | 1.42 | (510) |
| **Hypophosphatasia**<br>**(Patient 0124)** | 2.965 | 0.337268 | 3.37 | (002) |
|  | 3.28 | 0.304878 | 3.05 | (210) |
|  | 3.6325 | 0.275292 | 2.75 | (112) |
|  | 3.861 | 0.259 | 2.59 | (301) |
|  | 4.485 | 0.222965 | 2.23 | (221) |
|  | 4.6755 | 0.213881 | 2.14 | (311) |
|  | 4.887 | 0.204625 | 2.05 | (113) |
|  | 5.1695 | 0.193442 | 1.93 | (222) |
|  | 5.452 | 0.183419 | 1.83 | (213) |
|  | 5.8645 | 0.170518 | 1.71 | (004) |

**Supplementary Table S3:**
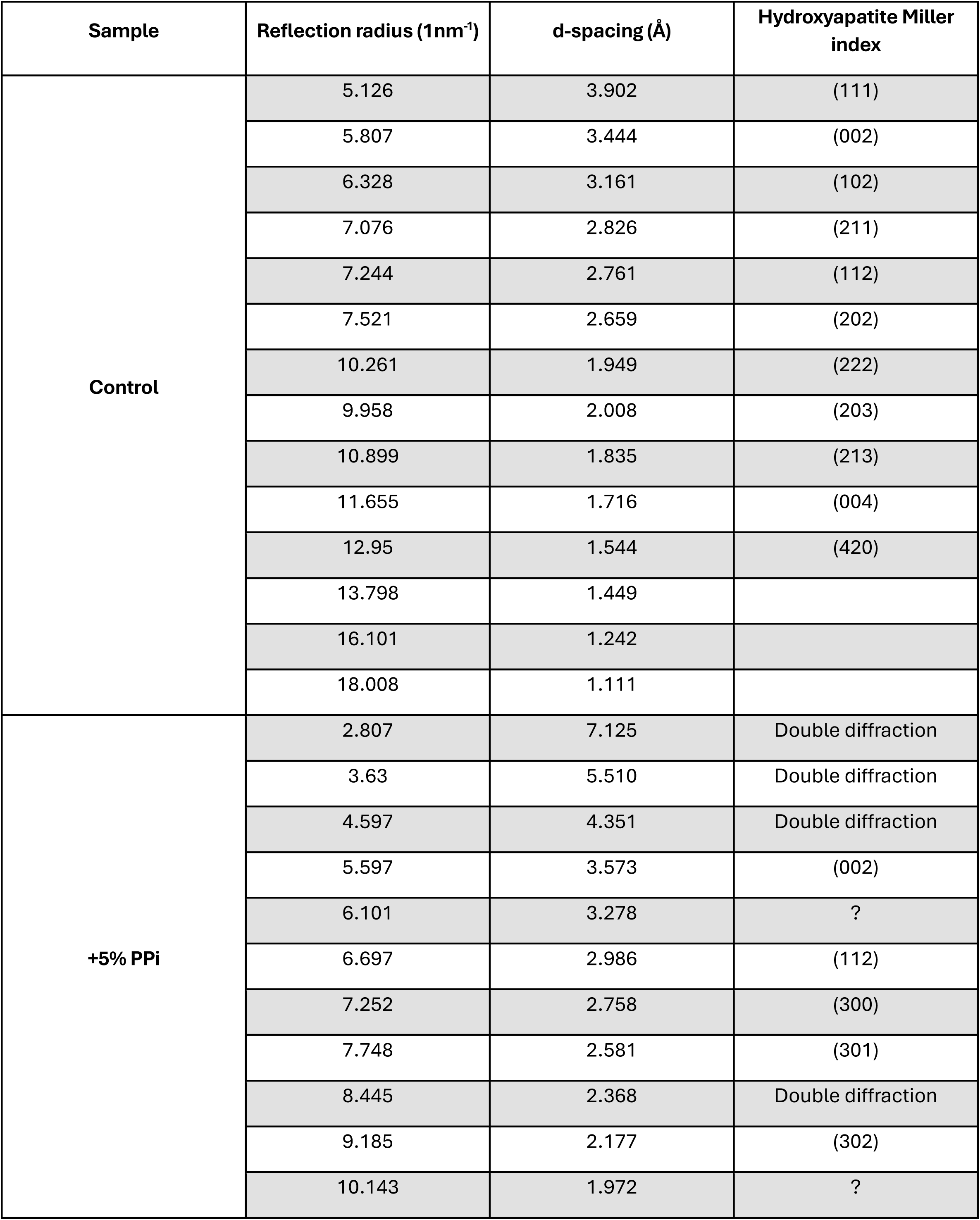
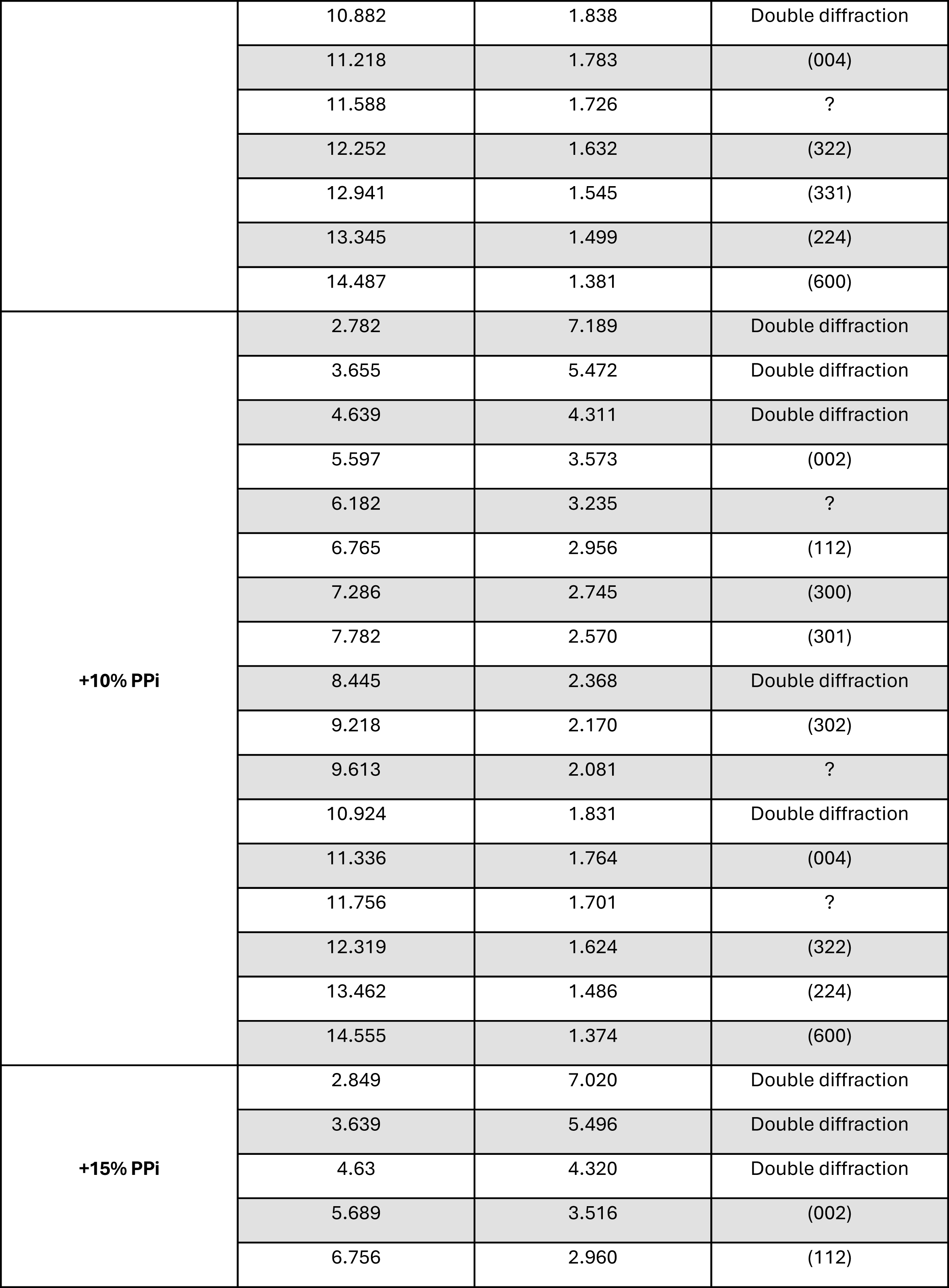
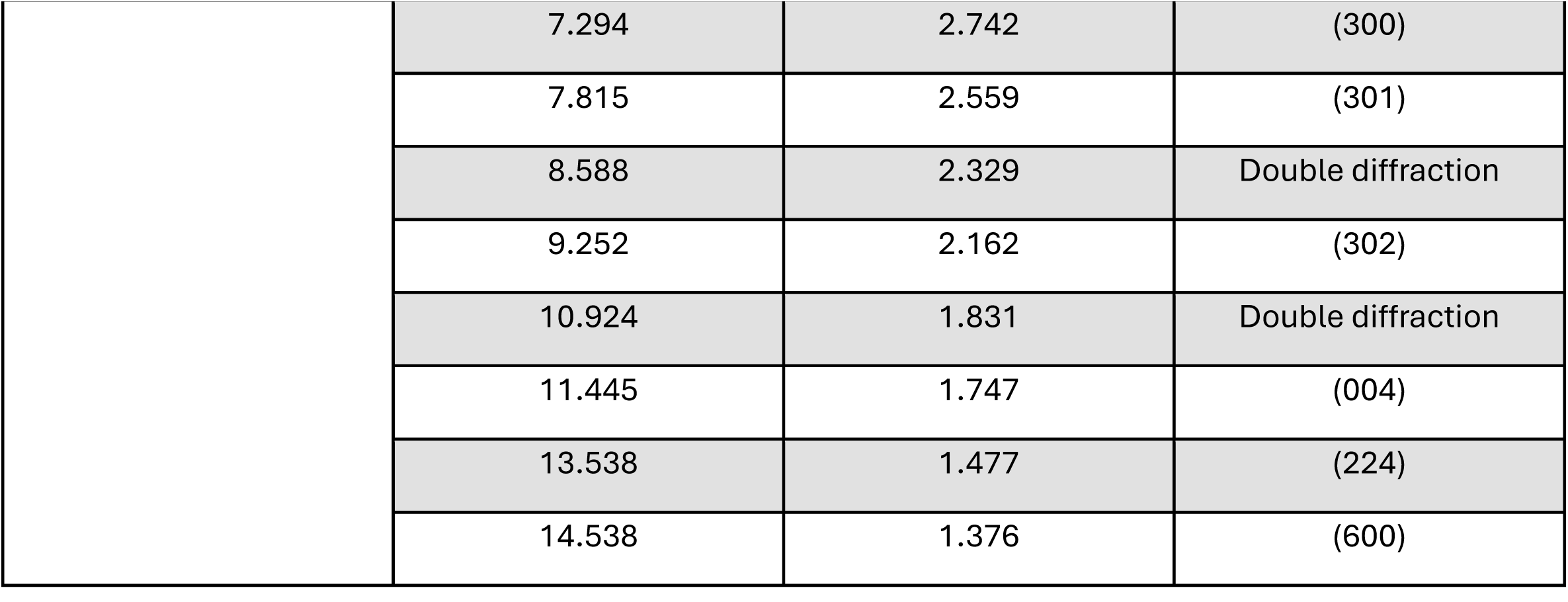
Full diffraction indexing of selected area electron diffraction (SAED) patterns obtained in synthetic bone mineral material samples. Hydroxyapatite Miller indices are assigned according to reference standard X-ray data in hydroxyapatite(*59*).

